# In-phase and in-antiphase connectivity in EEG

**DOI:** 10.1101/2021.05.19.444800

**Authors:** Christian O’Reilly, John D. Lewis, Rebecca J. Theilmann, Mayada Elsabbagh, Jeanne Townsend

**Affiliations:** Montreal Neurological Institute, Azrieli Centre for Autism Research, McGill University, Montreal, Canada; Montreal Neurological Institute, McGill University, Montreal, Quebec, Canada; Department of Radiology, UC San Diego, La Jolla, CA; Department of Neurosciences, UC San Diego, La Jolla, CA; Research on Aging and Development Laboratory, UC San Diego, La Jolla, CA

**Keywords:** zero-lag connectivity, functional connectivity, EEG, neural communication, coherency, interhemispheric connectivity

## Abstract

Zero-lag synchrony is generally discarded from functional connectivity studies to eliminate the confounding effect of volume conduction. Demonstrating genuine and significant unlagged synchronization between distant brain regions would indicate that most electroencephalography (EEG) connectivity studies neglect an important mechanism for neuronal communication. We previously demonstrated that local field potentials recorded intracranially tend to synchronize with no lag between homotopic brain regions. This synchrony occurs most frequently in antiphase, potentially supporting corpus callosal inhibition and interhemispheric rivalry. We are now extending our investigation to EEG. By comparing the coherency in a recorded and a surrogate dataset, we confirm the presence of a significant proportion of genuine zero-lag synchrony unlikely to be due to volume conduction or to recording reference artifacts. These results stress the necessity for integrating zero-lag synchrony in our understanding of neural communication and for disentangling volume conduction and zero-lag synchrony when estimating EEG sources and their functional connectivity.

## Introduction

The concept of functional connectivity provides a very useful construct to study brain dynamics in health and diseases. It is commonly estimated from hemodynamic responses recorded with functional magnetic resonance imaging (fMRI) or near-infrared spectroscopy (fNIRS). Alternatively, it can be measured from electrophysiological activity captured through electroencephalography (EEG) or magnetoencephalography (MEG). Although imaging based on hemodynamic responses supports a higher spatial resolution, its relationship with the underlying neural activity is indirect. Further, the electrophysiological activity occurs on a much faster time scale than the comparatively slow hemodynamic response. For that reason, MEG and EEG (M/EEG) provide a much higher temporal precision and allow for the study of neuronal oscillatory patterns and their correlates. However, the estimation of functional connectivity from M/EEG signals is associated with many pitfalls. M/EEG can only be recorded when synchronous discharges of spatially aligned populations of neurons, like the pyramidal cells, generate a cumulative magnetic/electric field that propagates by volume conduction from its source all the way to the sensors, e.g., surface electrodes in EEG or superconducting quantum interference devices (SQUID) in MEG. Unfortunately, the same volume conduction that makes M/EEG possible has been shown to be a major confounder for functional connectivity^1^. To overcome this limitation, neuronal communication based on action potentials needs to be distinguished from spurious dependencies due to volume conduction. Since the electromagnetic propagation can be considered instantaneous in the context of M/EEG^2^, volume conduction is sufficient, but not necessary, for in-phase or in-antiphase M/EEG activity between distant locations on the head. Zero-lag synchronization has consequently often been purposefully discarded ^3–6^. Such approaches reject any unlagged synchronization contributing to the estimates of functional connectivity, regardless of whether it is an artifact of volume conduction or the result of genuine functional connectivity synchronized with no lag. Although counter-intuitive, synchronization between oscillatory systems linked through delayed connections (i.e., as is the case for the delayed communication between brain regions due to the finite propagation speed of action potentials along the axons) is possible in systems like biological neural networks because of the presence of feedback loops. Although much less studied than phase-locking, evidence for the existence of zero-lag neuronal connectivity has been put forward in fundamental^7^, experimental^8–13^, and modeling studies^14–16^.

In a recent study^12^, we demonstrated using intracranial recordings — a recording modality not significantly sensitive to volume conduction above ∼2 cm^1, 17, 18^ — in presurgical patients with drug-resistant epilepsy that zero-lag connectivity was common between long-distance (> 10 cm) homotopic brain regions. Further, these analyses demonstrated that this interhemispheric synchrony was three times more likely to be in-antiphase than in-phase, a property interesting because it might relate to interhemispheric inhibition^19–21^ supported by the corpus callosum and shown to be involved in motor control^22–24^, lateralized overt attention^25, 26^, and somatosensory processing^27, 28^. These observations also provide a very specific pattern of synchrony that can be leveraged to distinguish the presence of genuine zero-lag connectivity from volume conduction in other modalities such as EEG.

Although this previous intracranial study provided convincing evidence that zero-lag connectivity is common in normal interhemispheric brain dynamics, the extent to which it plays an important role in M/EEG functional connectivity needs to be confirmed. Due to the greater proximity between the sensors and the neuronal activity as well as the smaller size of the sensors, local field potentials (LFP) recorded in intracranial recordings can resolve patterns of activity on a much smaller spatial scale than M/EEG. Thus, although the importance of zero-lag synchronization between distant cell assemblies has been demonstrated in LFP, it is yet unclear if such synchronization is happening within spatial scales contributing to M/EEG.

Further, various additional factors are involved in making such a synchronization measurable or not in M/EEG. For example, if these patterns of synchrony are supported by cell assemblies comprising not enough cells or if the electromagnetic dipolar sources generated by these cells are not sufficiently aligned, they may create an electromagnetic field that has a significant local amplitude but that is mostly closed on itself and therefore not visible from the outside of the brain. However, if the presence of zero-lag connectivity can be demonstrated in M/EEG, this would imply that methods to estimate functional connectivity need to be revised to control for the confounding effect of volume conduction without discarding altogether all zero-lag interactions between M/EEG channels.

To answer these questions, we analyzed EEG recordings from healthy participants. First, we tested the hypothesis, based on previous LFP evidence, that EEG zero-lag connectivity will be dominated by antiphase synchrony specifically along the interhemispheric (IH) axis. We further demonstrated through a set of simulations that this effect could not be ascribed only to volume conduction. Second, we performed a more exploratory and descriptive analysis to characterize how zero-lag connectivity is distributed in the brain and its relationship to spatial and temporal frequencies of EEG recordings. Third, we examined the potential confound of using an average reference in producing false zero-lag synchrony, i.e., by definition EEG reference schemes relying on a common reference subtract the same reference signal to all channels and can therefore add a non-lagged component across the channels. We estimated the reference-free scalp current density (SCD) and demonstrated that estimates of zero-lag connectivity remained reliable when using a reference average.

## Results

### The predominance of in-antiphase synchrony in the interhemispheric (IH) direction

We first analyzed the phase of coherency between pairs of channels in EEG recordings from 63 healthy adults (see methods for detail). Figure 1.a-c shows coherency for an example subject and for a given pair of channels (PO7 and PO8). For panels b and c, the coherence and the phase of the coherency have been computed over 1000 randomly selected epoch sub-samples (i.e., bootstrapping). The grayscale profile represents the density of the estimated coherency and supports the stability of these estimates. As depicted in this example, the phase was most often stable at either 0 or π rad across frequencies, with sometimes more variability around the alpha/beta frequency range. In-antiphase synchrony was very clearly dominating for IH pairs, whereas the opposite was evident for anteroposterior (AP) pairs, as shown in Figure 1.d-g. To validate this observation across subjects, we selected two pairs centered around the Cz channel, one along the IH direction (C3-C4) and the other along the AP (Pz-Fz) direction. Again, the contrast, illustrated in Figure 1.h,i is startling, with 25 out of 63 subjects showing predominantly in-antiphase synchrony (π/2 < ϕ < 3π/2for all frequencies) for the IH pair and none showing in-phase synchrony on the same pair. The opposite pattern is observed on the AP pair, with 11 out of 63 subjects showing predominantly in-phase activity (− π/2 < ϕ < π/2 for all frequencies) for this pair of channels and none showing in-antiphase synchrony. A similar analysis was performed for all IH and AP pairs for every subject and we plotted in Figure 1.j the distribution across subjects of the proportion of pairs that were synchronized in-antiphase, in-phase, or neither (undetermined). The larger amount of in-phase synchrony is evident for AP pairs (t-test: t=-26.608; p=4.30e-53; mean ± std: IH=0.196 ± 0.066; AP=0.681 ± 0.129; N=63; Cohen’s d=-3.352) as is evident the larger number of in-antiphase synchrony for IH pairs (t-test: t=24.983; p=3.15e-50; mean ± std: IH=0.437 ± 0.132; AP=0.010 ± 0.029; N=63; d=3.148). Aside from the very small p-values obtained for these t-tests, the corresponding effect sizes are quite large compared to those typically observed in EEG and functional connectivity studies given that Cohen’s d is generally considered to reflect a “large” effect size for values close to 0.8^29^ and a “huge” effect size for values close to 2.0^30^. Types of synchronization (in-phase versus in-antiphase) are also illustrated for 10 randomly selected participants for IH (Figure 1.k) and AP (Figure 1.l) pairs, further demonstrating how clear and stable across subjects this relationship is.

**Figure 1.**
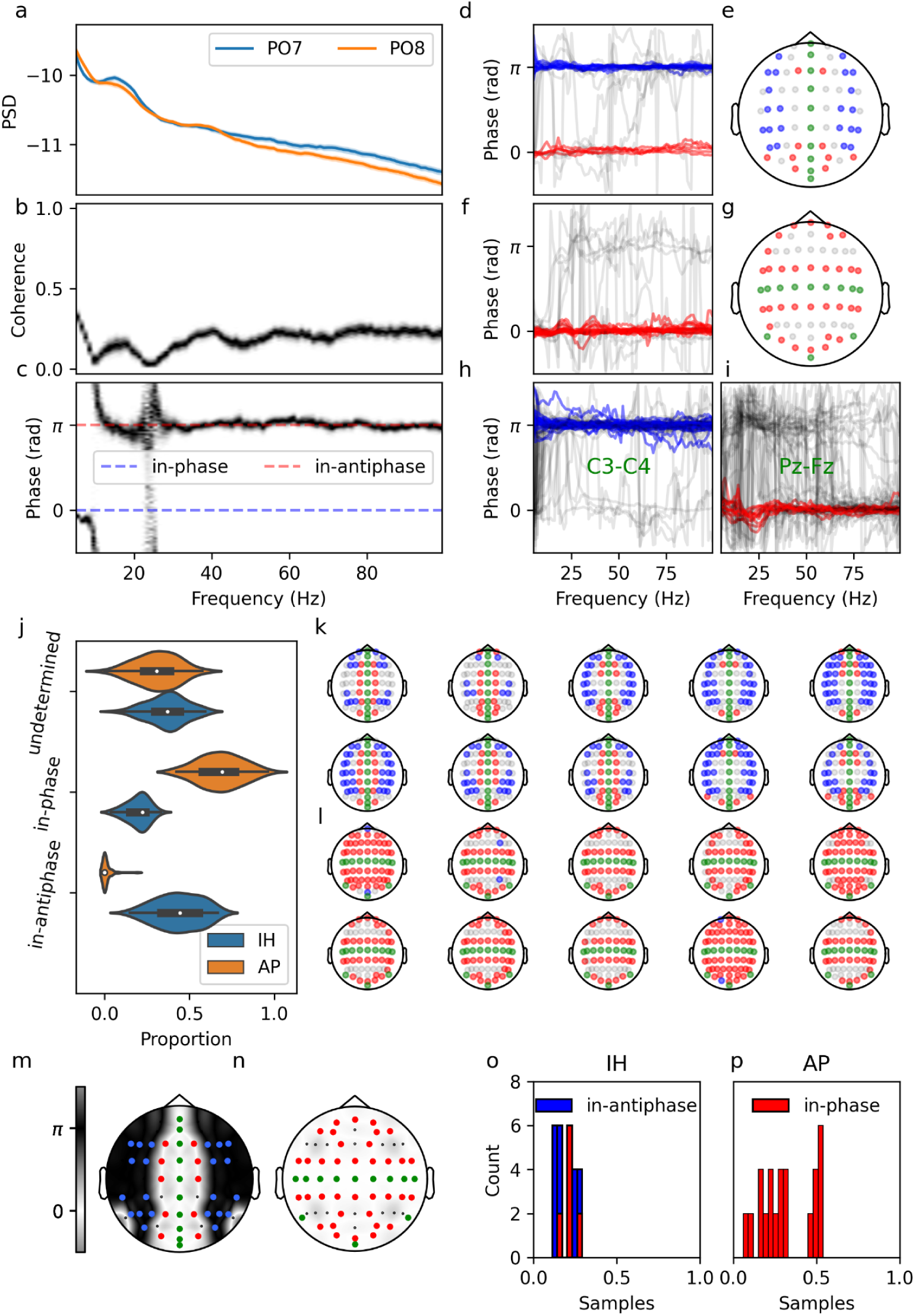
Comparative analysis of IH and AP channel pairs from EEG recording. Panels a-g are for an example subject; panels h-j and m-p are across subjects. a-c) Example of power spectral density (PSD; a), coherence (b), and phase of coherency (c) for a given channel pair (PO7, PO8). d) Phase for IH pairs, color-coded with pairs predominantly in-phase (red), in-antiphase (blue), and or neither (undermined; gray). e) Topomap showing the positions of the channels used for the time-series plotted in panel d, with the same color-coding except for green which is used for unpaired channels. f,g) Similar to d,e but for AP pairs. h,i) Phase of coherency between C3-C4 (IH; h) and Pz-Fz (AP; i) for all subjects (one line per subject), color-coded as described for panel d. j) Distribution across subjects of the proportion of in-phase, in-antiphase, and undetermined pairs. k,l) Synchronizations for IH (k) and AP (l) pairs for ten randomly selected subjects. m,n) Average phase across subjects for the IH (m) and AP (n) pairs. Red and blue dots are indicating sensors where the average phase was less than one time sample away (see text) from 0 (in-phase; red) or π (in-antiphase; blue) rad. o, p) Distribution of the difference between the mean phase and 0 or π (in the corresponding number of time samples) for the set of in-phase and in-antiphase pairs identified in panel n,m and for the IH (o) and AP (p) directions.

We further computed the average phase across subjects over the 30-100 Hz band (to exclude lower frequencies showing increased variability) for IH (Figure 1.m) and AP (Figure 1.n) directions. Again, the contrasting pattern is very clear. The overlaid colored sensors show pairs which 95% interval of confidence (IC) on their mean phase included 0 (in-phase; red) or π (in-antiphase; blue) and that were at most including a deviation of one time sample (i.e., <3.9 ms) away from these values. More specifically, since converting phases into time intervals can only be done at a specific frequency, 3.9 ms corresponds to 0.736 rad at 30 Hz and 2.454 rad at 100 Hz. We took the most restrictive criterion (i.e., 0.736) and we determined the phase within the 95% IC of the mean (across subject) that was the farthest away from either 0 or π and divided this value by 0.736 to assess how many time samples it was away from 0 or π. For example, if the 95% IC on the mean of the phase for the O1-O2 pair is [-0.3, 0.1], then it is considered to be at most 0.3/0.736=0.408 time samples aways from 0 (in-phase). Note that since we take the most extreme point of the IC and use the criterion that was the most severe, this is a rather conservative definition. The distribution of these departures from 0 or π rad for the pairs highlighted in panel m,n shows that they are tightly synchronized with no significant lag, if any (Figure 1.o,p).

### Volume conduction introduces no directional bias in zero-lag connectivity

We performed an additional analysis on simulated and surrogate signals to rule out the possibility that the previous results were due to directional biases in how volume conduction travels or in the orientation of the dipolar sources. For example, since dipolar sources tend to produce anti-phased electrical fields in opposite directions along their axis, the size of the interhemispheric fissure could be hypothesized to favor in-antiphase synchronization in the IH direction (i.e., dipoles in interhemispheric fissures are mostly aligned with the IH direction and could therefore produce anti-phased fields on each side of the head). Thus, we simulated scalp signals from the estimated sources and analyzed them in a similar fashion as we did for the recorded EEG signals, with (surrogate) and without (simulated) randomly shuffling the position of the sources within the cortex before simulating the EEG generated from these sources. Since the simulated EEG closely reproduces the recorded EEG (see Figure 10 in online methods), the results from the coherence analysis using the simulated signals are very similar to those reported for the recorded EEG in Figure 1 (see Supp. Figure 1). We observe more in-phase synchronisation along the AP direction (t=-27.356; p=2.27e-54; mean ± std: IH=0.177 ± 0.070; AP=0.714 ± 0.139; N=63; d=-3.447) and more in-antiphase synchronisation along the IH direction (t=29.066; p=3.38e-57; mean ± std: IH=0.516 ± 0.135; AP=0.009 ± 0.031; N=63; d=3.662).

By contrast, when using the surrogate dataset to assess the effect of volume conduction without any systematic underlying spatial pattern in the distribution of the cortical sources, we observed no statistical difference in the proportion of in-phase (t=-0.35; p=0.72332; mean ±std: IH=0.34 ± 0.12; AP=0.34 ± 0.11; N=100; d=-0.035) or in-antiphase (t-test: t=-0.34; p=0.73465; mean ± std: IH=0.35 ± 0.13; AP=0.35 ± 0.15; N=100; d=-0.034) pairs using a set of 100 randomization within our example subject (Figure 2.a-j). Similarly, we observed no difference in the proportion of in-phase (t-test: t=1.47; p=0.14545; mean ± std: IH=0.34 ± 0.13; AP=0.31 ± 0.10; N=63; d=0.185), but potentially a small difference (possibly a false positive) in the proportion of in-antiphase (t-test: t=-2.04; p=0.04383; mean ± std: IH=0.34 ± 0.13; AP=0.39 ± 0.14; N=63; d=-0.257) pairs across subjects (Figure 2.k-m). These observations further confirm that our results showing significantly larger in-antiphase synchrony in the IH direction between homotopic channel pairs in recorded EEG cannot be attributed to a bias in the directionality of dipolar sources or of volume conduction due to structural aspects of head tissues.

**Figure 2.**
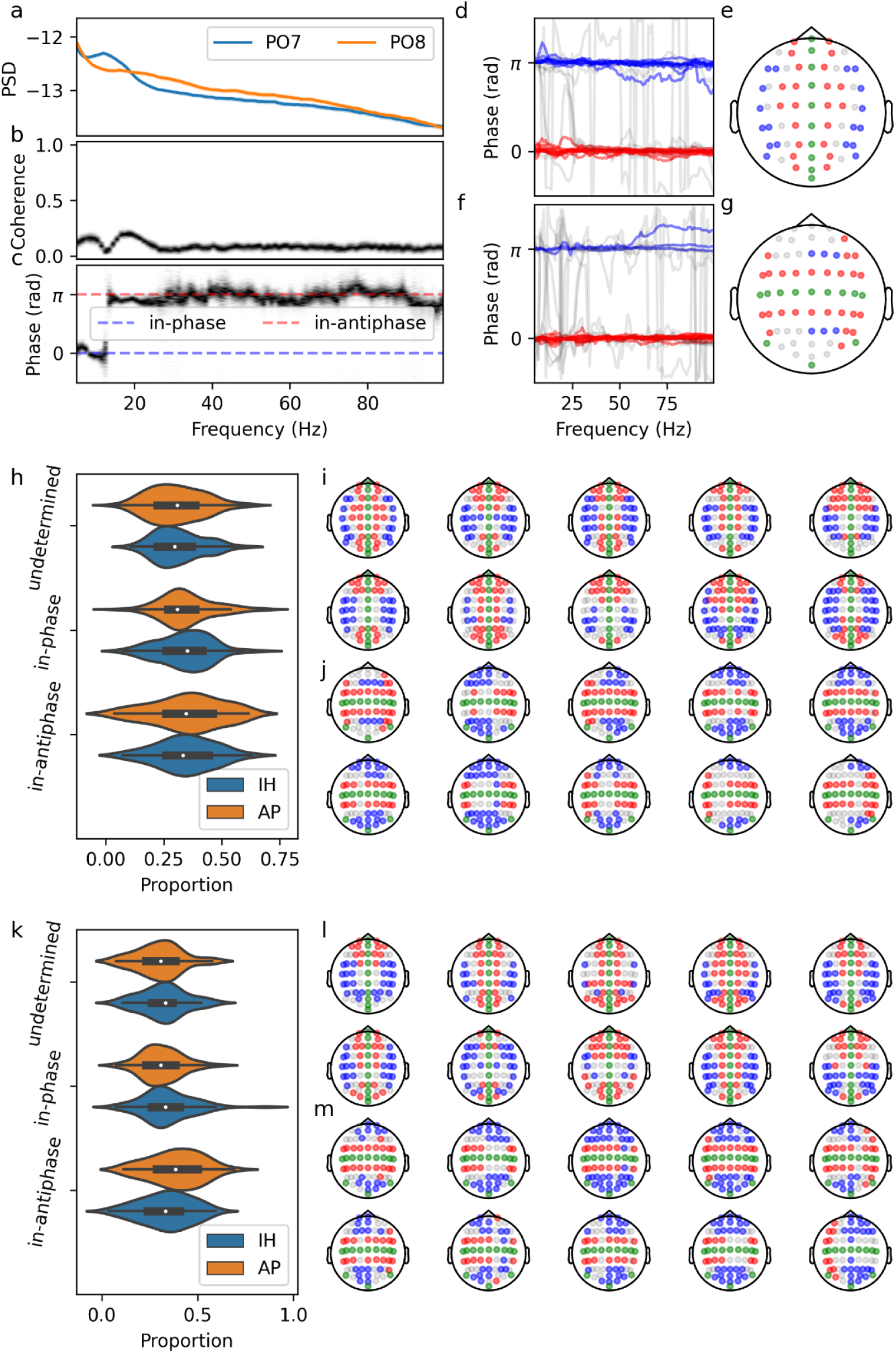
Comparative analysis of IH and AP channel pairs for a surrogate dataset obtained by simulating EEG from spatially randomized cortical sources. Panels a-j are computed for an example subject (same as in Figure 1), while panels k-m are showing results across subjects. a-c) Example of PSD (a), coherence (b), and phase of the coherency (c) for the channel pair (PO7, PO8) on an example subject. d-g) These panels show the same data as for Figure 1.d-g, but for the surrogate EEG. h) Distribution of the number of pairs per synchronization type, across 100 randomizations. i,j) Color-coded dominant phase (blue: in-antiphase; red: in-phase; grey: undetermined; green: unpaired channels) for 10 different randomizations, for the IH (i) and the AP (j) directions. k-m) Same as h-j, but across subjects.

### Zero-lag directional patterns using scalp current density (SCD)

Having obtained clear corroboration of our hypothesis about the predominance of in-antiphase synchronization in the IH direction, we performed a second analysis to 1) rule out the potential confounding effect of using an average reference, 2) to explore the effect of frequency, and 3) to investigate more thoroughly the spatial distribution of the zero-lag connectivity. We first computed the SCD from recorded and surrogate EEG to get rid of the impact of the reference and we plotted in Figure 3 polar heatmaps illustrating the proportion of subjects with in-phase or in-antiphase synchrony for the different pair orientations and inter-electrode distances, using pairs as defined in Figure 9.c (online methods). On these plots, we can observe that zero-lag synchrony, as opposed to volume conduction, depends on the frequency. We also note that spatial patterns of in-phase and in-antiphase synchrony appear not to be as clear in this SCD analysis as in the previous average reference analysis, although the distribution of zero-lag synchrony with respect to the orientation of the channel pairs is still clearly not random or uniform (e.g., in-phase at 10 Hz). The proportion of pairs that were shown to be above the 99% IC of their surrogate counterpart at 10, 20, 40, and 70 Hz was, respectively, 22.8%, 22.3%, 16.8%, and 17.5% for the in-phase synchrony and 8.39%, 5.27%, 9.11%, and 10.6% for the in-antiphase synchrony, all well above the 1% chance level.

**Figure 3.**
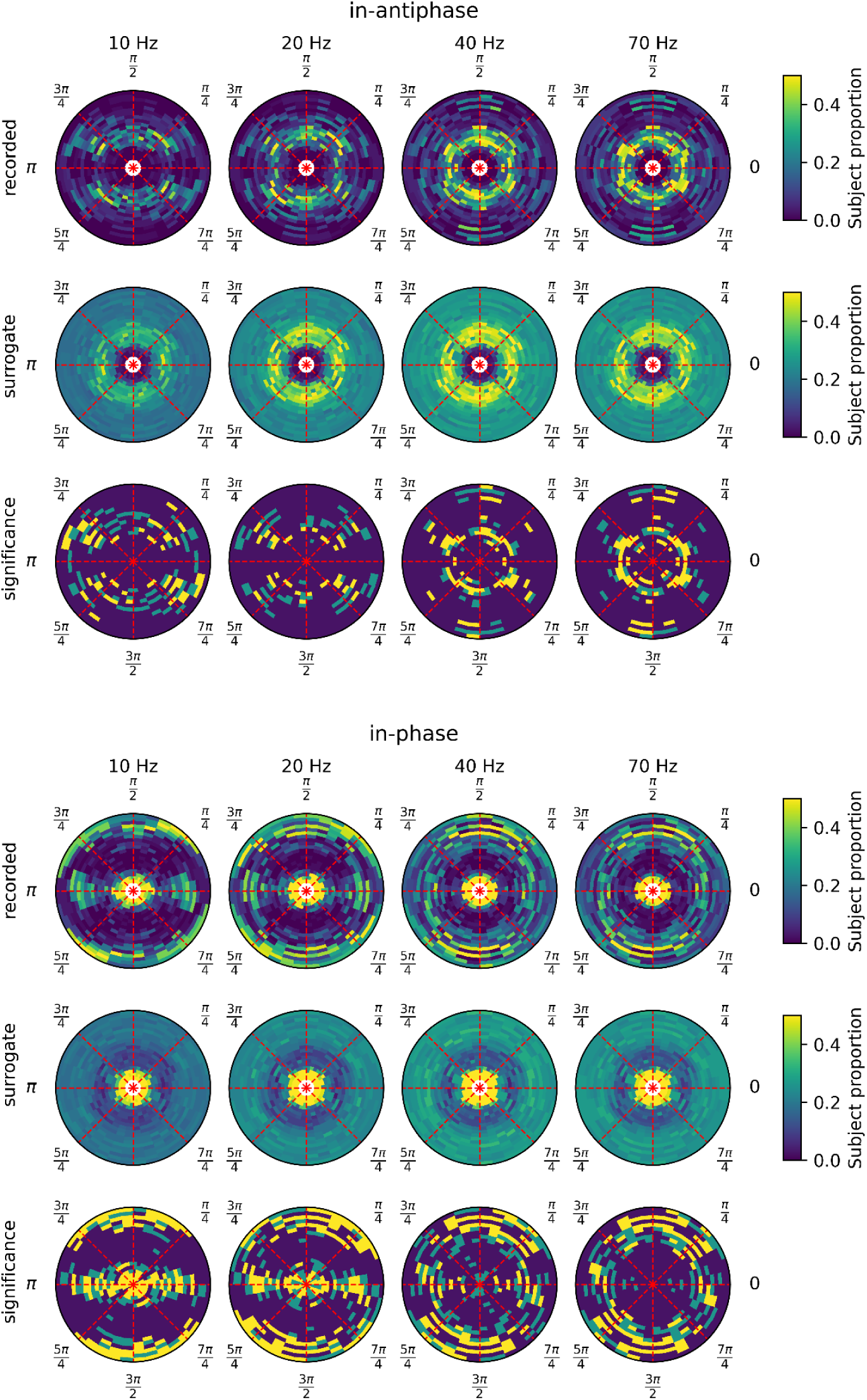
Polar plots showing the proportion of subjects with in-antiphase (upper panel) or in-phase (bottom panel) synchrony as a function of the channel pair orientation (angular coordinate) and the inter-electrode distance (r coordinate). Each column shows the results for a different frequency (10 Hz, 20 Hz, 40 Hz, and 70 Hz). On the polar plots, a 0 rad angle corresponds to IH, whereas π/2 rad corresponds to AP. The first row of each panel gives the values from the recording, the second row shows the 99th percentile of the surrogate distribution, and the third row shows the statistical significance (dark blue: not significant; light blue: significant at p<0.01, not corrected for multiple tests; yellow: significant at p<0.01, Bonferroni-corrected for testing independently 834 channel pairs).

### Impact of the frequency on zero-lag connectivity

In order to get a more complete picture of how the phase of coherency varies across frequencies and channels, we sorted channel pairs by inter-electrode distance and plotted their average phase (across subject) as a function of the frequency (Figure 4). Although this heatmap makes it clear that the phase-frequency relationship is strongly impacted by the inter-electrode distance, it also shows significant variability for any given distance. To identify possible patterns of functional connectivity, we clustered channel pairs depending on their phase-frequency dynamics. Although these clusters have different topological properties (e.g., compare the lateral AP connections in the second topomap from the top with the frontal and posterior IH connections in the second topomap from the bottom), their patterns are rather complex and not easily synthesized as clear organizational rules.

**Figure 4.**
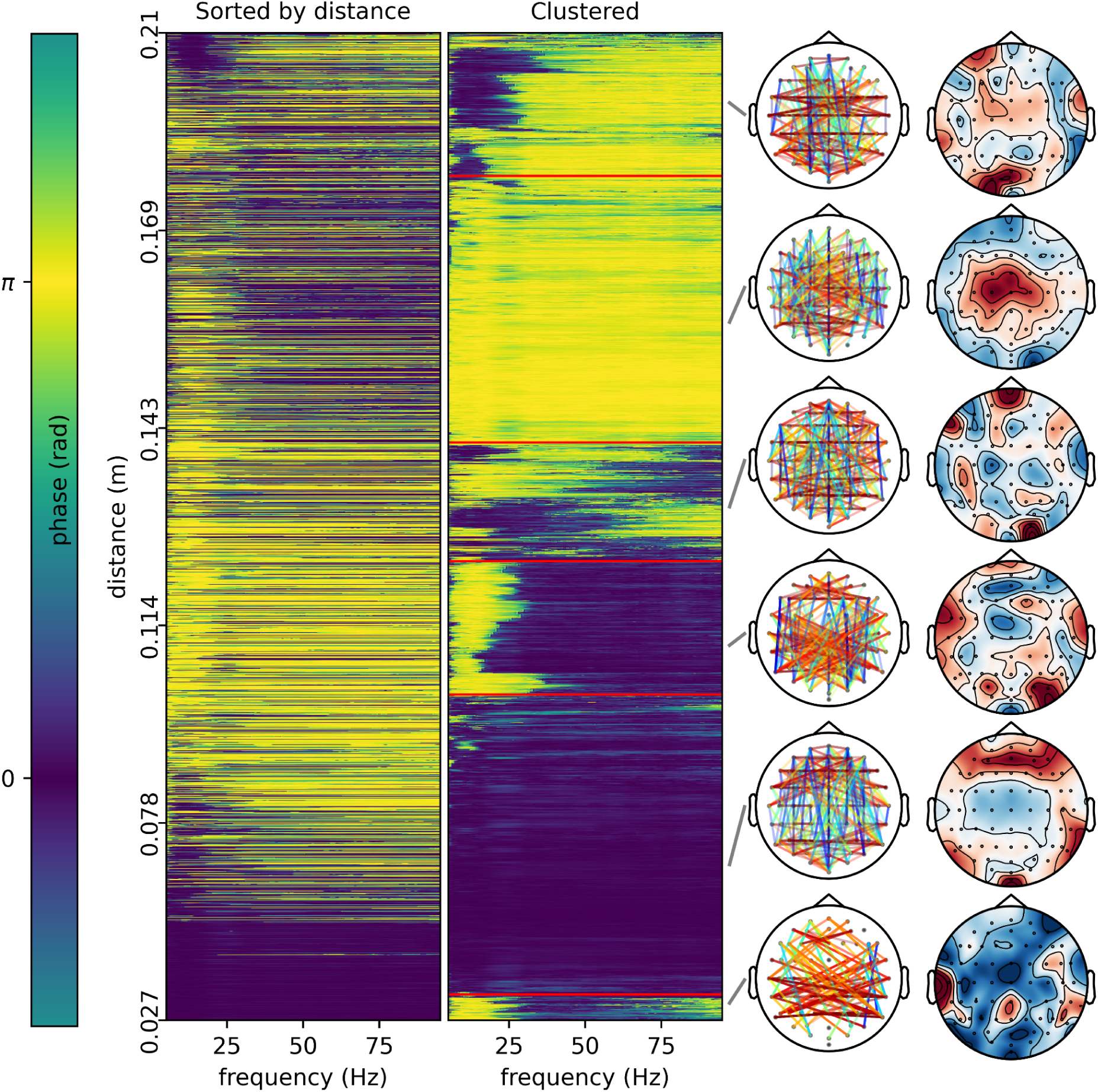
Phase of the coherency as a function of the frequency. On the first panel, the electrode pairs are sorted by increasing inter-electrode distance. Note that the y-axis in this plot is not linear with respect to inter-electrode distance because the density of channel pairs varies as a function of the distance (see Supp. Figure 3 for the distribution of the inter-electrode distances). In the second panel, channel pairs have been reordered using hierarchical clustering to group them by similarity. These pairs have been split (red horizontal line) into six groups that were subjectively considered mostly homogeneous. Topomaps showing the connections (up to a maximum of 200 randomly sampled pairs to avoid overcrowding the graphs) for these six clusters are drawn on the side. The connections are color-coded depending on their orientation (IH: red, AP: blue; intermediate values following a rainbow palette) to further help identify differences in general orientation between clusters. The rightmost topomaps show the relative proportion of connections made by the different channels.

### Impact of distance on zero lag-connectivity

To help further unpack the topological variations we observed, we plotted the proportion of subjects showing a given type of synchrony (in-phase, in-antiphase, or lagged) for each channel pair, as a function of the inter-electrode distance (Figure 5), for the same four frequencies used in Figure 3. We overlaid on these scatter plots the 99th percentile of the surrogate distributions (grey dots). Then, using a sliding window, we computed a moving 99th percentile (i.e., a 99th percentile across channel pairs within the window, the value for each included pair being already estimated as the 99th percentile of their surrogate distribution) to establish a significance threshold (shown as a red dashed line) that depends on the inter-electrode distance. Using this threshold as well as 99th percentile for the individual pairs (i.e., for each pair, both criteria must be met), we selected channel pairs for the in-phase, in-antiphase, and lagged type of synchrony and displayed these pairs on topomaps. We note that the general relationship between the distance and the types of synchrony for the surrogate data is coherent with what can be expected from volume conductions for dipolar sources: large in-phase synchrony at short distances shifting to in-antiphase synchrony at larger distances, with a level of lagged synchrony being rather stable after the initial low proportion at short distances due to the dominant effect of volume conduction.

**Figure 5.**
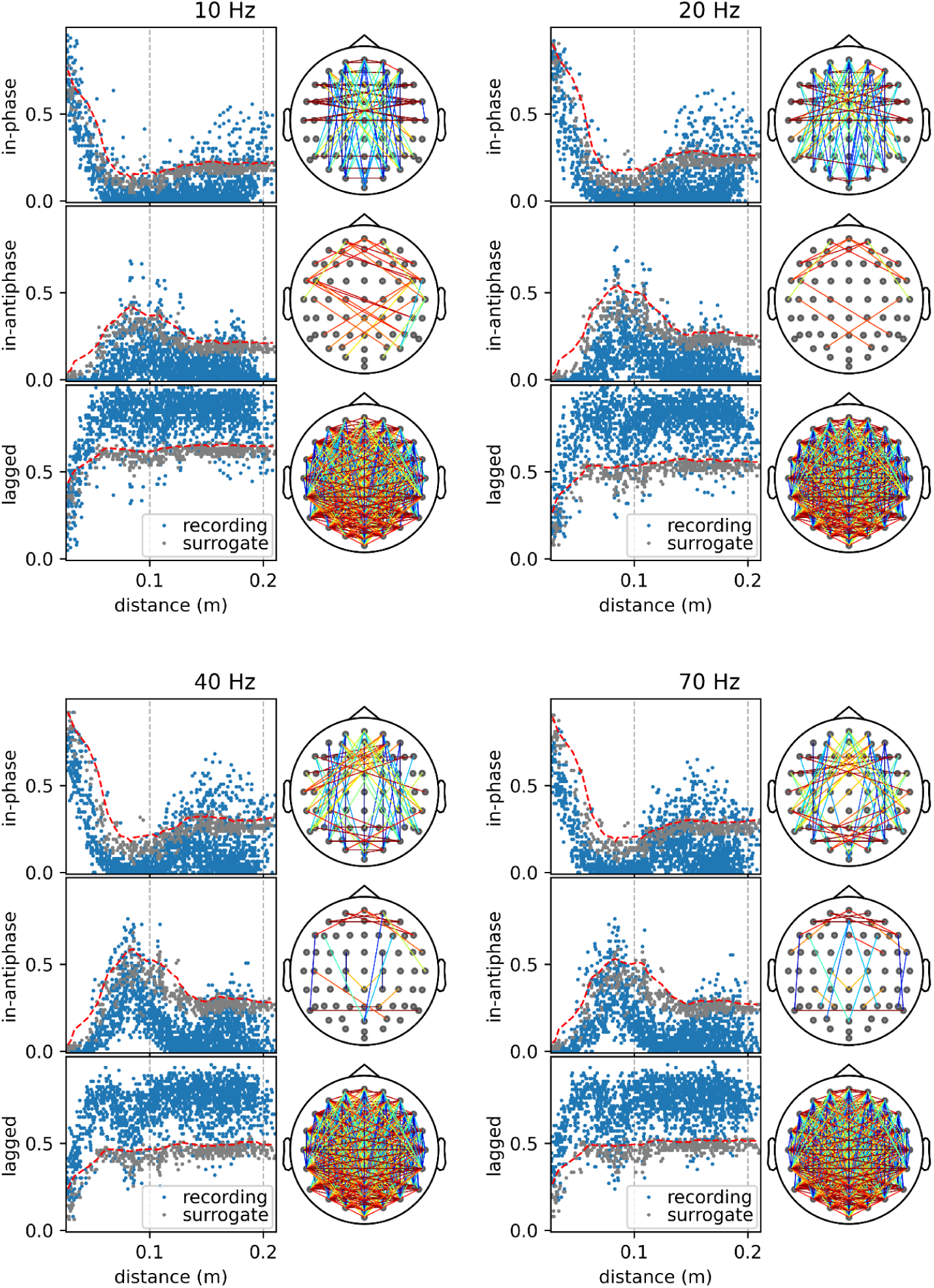
Selecting channel pairs based on distance-dependent thresholds. Four panels present the same analysis at 10, 20, 40, and 70 Hz. Each panel has three rows for in-phase, in-antiphase, and lagged synchrony. Each row contains a scatter plot and a topomap. The scatter plot shows dots associated with recording EEG (blue) and the 99th percentile of the surrogate EEG (grey). Each dot reports the proportion of subjects showing this type of synchrony for a specific pair. We computed a 99th percentile of the cloud of surrogate points using a sliding window to obtain a smooth distance-dependent threshold (red dashed lines). Pairs that had a higher proportion of subjects than the 99th percentile of their corresponding surrogate distribution and that were above this distance-dependent threshold were considered statistically significant and were displayed in the topomaps. The connections in the topomaps are color-coded with respect to their orientation (IH: red, AP: blue, intermediate values following a rainbow palette).

### The predominance of in-antiphase synchrony in the IH direction is not due to common reference artifacts

To evaluate more systematically the relationship between the type of synchrony and the orientation of the channel pairs, we used an orientation index corresponding to a weighted average of 2D points (amplitude, orientation) where the amplitude is weighted by a cosine function peaking at every π multiple of an angle θ (see online methods for the complete description). For this analysis, all pairs were used and their amplitude was defined as a boolean value equal to one only for the statistically significant pairs (i.e., the pairs displayed on the corresponding topomaps of Figure 5). In short, this index is computed for every subject and every value of θ within [-π/4, 3π/4]. It peaks at the angle that most selected electrode pairs are aligned with. As can be seen in Figure 6, the preferred angle for in-antiphase synchrony tends to reverse from AP at low frequencies to IH at high frequencies. This index highlights a clear and statistically significant increase of in-antiphase synchrony along the IH axis. Since this observation was made using SCD (reference-independent), these results validate that our previous observations concerning the predominance of in-antiphase synchrony along the IH direction were not due to artifacts introduced by the use of a common reference scheme.

**Figure 6.**
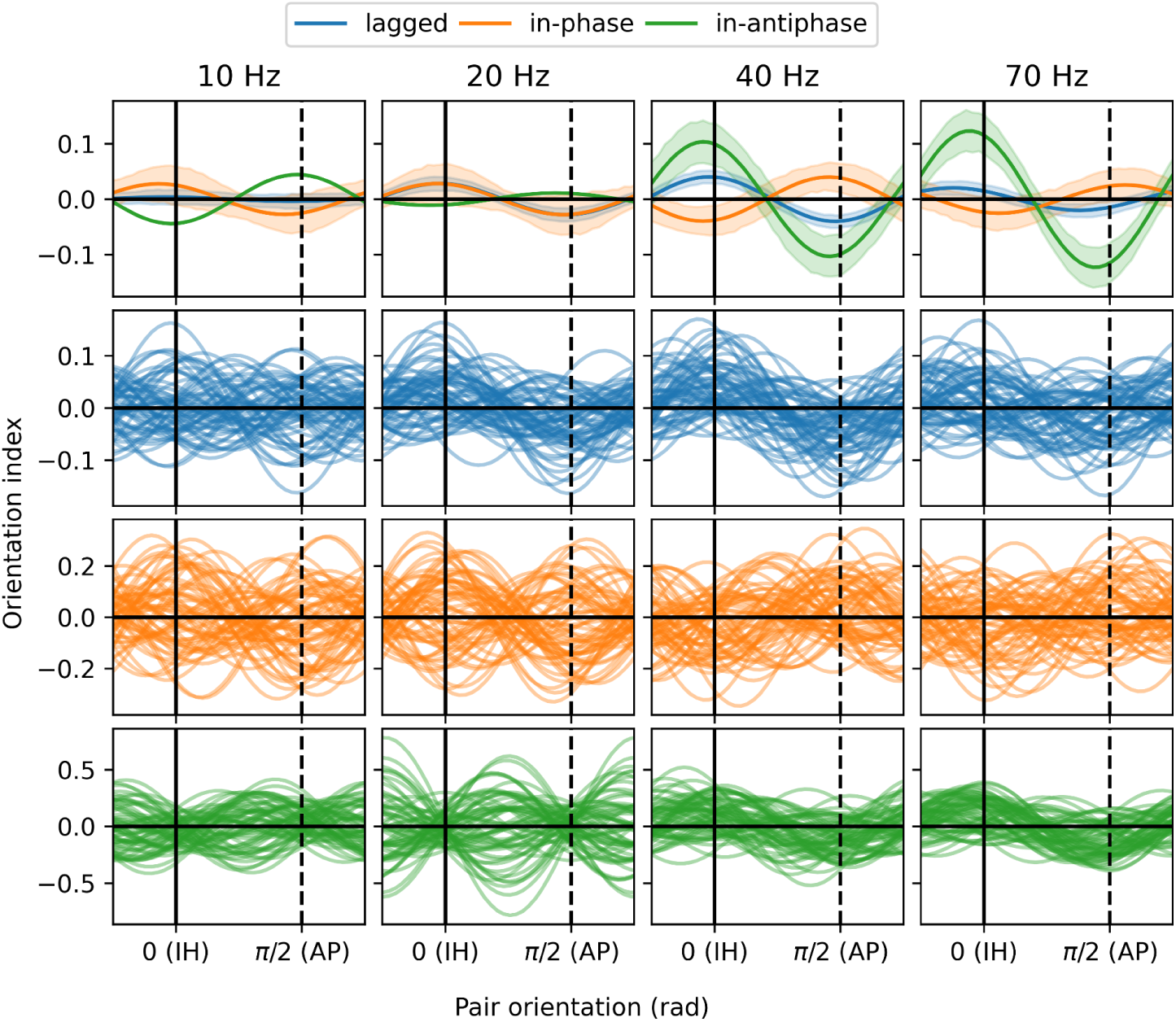
Orientation index for in-phase, in-antiphase, and lagged synchrony. The upper row shows its median value and its 95% IC across subjects, whereas the three other rows show the individual subjects.

### The complex patterns of zero-lag connectivity cannot be accounted for simply by volume conduction

Finally, to show a complementary and comprehensive picture of how in-phase versus in-antiphase synchrony varies across the scalp, we used, in turn, different channels as probes to display how zero-lag synchrony varies between these probes and the other electrodes (Figure 7). To efficiently show the patterns of in-phase versus in-antiphase synchrony, we used another index (the zero-lag index; see online methods) that takes values close to 1 when the distribution of the phase is mostly aligned with 0 (in-phase), close to -1 when this distribution is closely aligned with π (in-antiphase), and converge toward 0 when this distribution concentrates at neither of these two values. We plotted the variation of this index for the recorded EEG, the mean value of the surrogate distribution, the difference between the recorded and mean surrogate EEG, and the level of statistical significance (signed and in log_10_ scale) under the assumption that the surrogate distribution is normally distributed (Figure 7). The distribution for the surrogate data follows closely what we expect from volume conduction, which is a high level of in-phase synchrony at short distances decreasing regularly with distance in all directions and reversing toward in-antiphase synchrony (due to dipoles tangential to the scalp) at larger distances. The patterns that we see on the recordings, however, are much more complex and that is clearly reflected in the pattern of statistical significance. It would be hard, given these results, to consider that the zero-lag synchronization does not have a very significant component that is unrelated to volume conduction.

**Figure 7.**
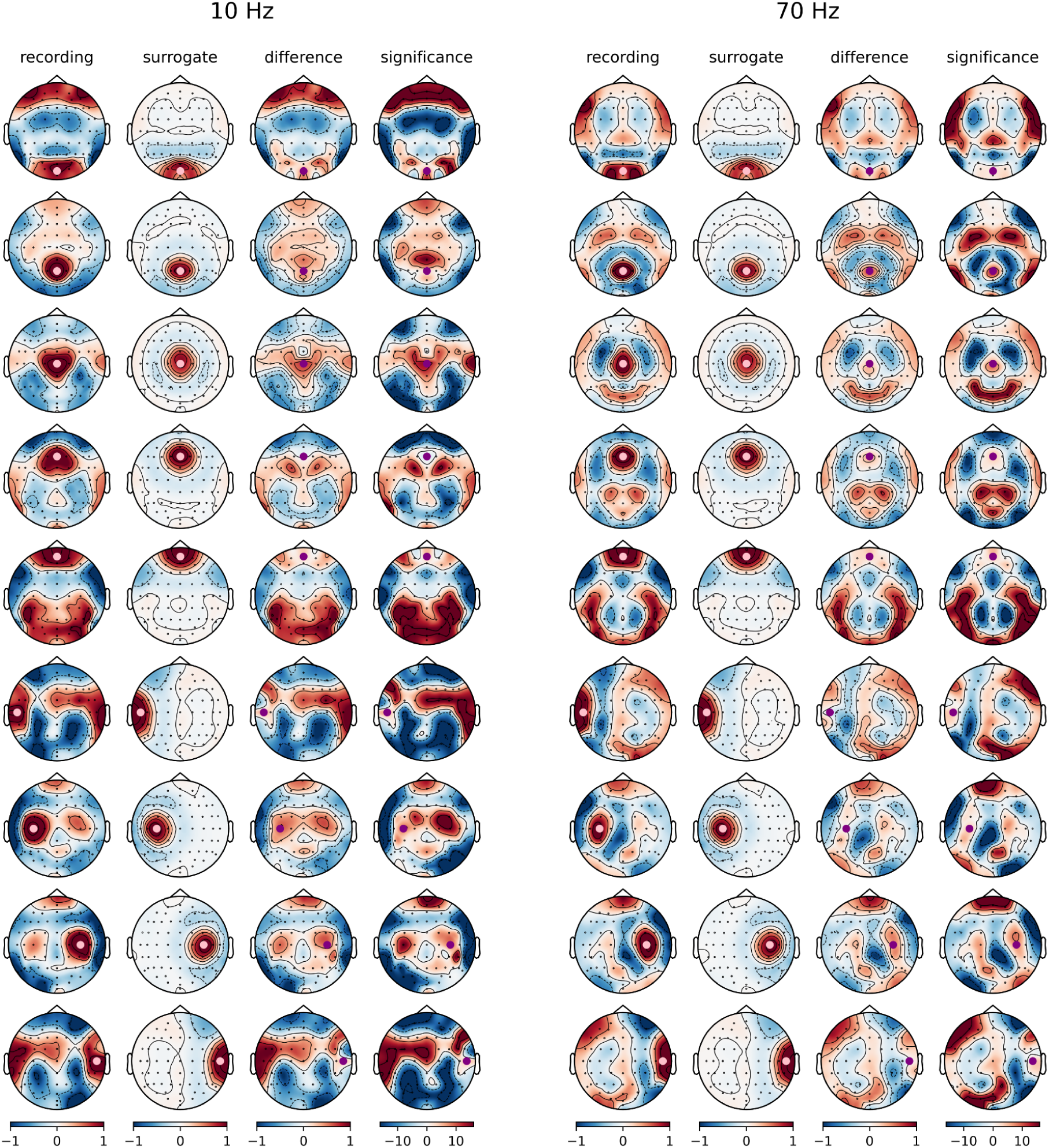
Patterns of variation for the zero-lag index (1: in-phase, -1:in-antiphase, 0: neither) using various probe electrodes (from top to bottom: Oz, Pz, Cz, Fz, Fpz, T7, C3, C4, T8; shown as a pink/purple circles) at 10 Hz (left) and 70 Hz (right). The first two columns show the topography of the zero-lag index computed on the recorded and the mean surrogate EEG, respectively. The third column shows the difference between the two first columns and the fourth is the log_10_ p-values under the null hypothesis that the zero-lag index is zero and that the surrogate values are normally distributed. These log_10_ p-values have been signed positive for in-phase and negative for in-antiphase synchrony. Reported log_10_ p-values are not corrected for multiple tests. Thus, the 0.05 statistical significance threshold adjusted using a Bonferroni correction for 64 tests on a log_10_ scale corresponds to absolute values greater than 3.1 on these plots.

### Zero-lag synchrony is most significant at lower spatial scales

Both the average reference and the SCD analyses clearly support the existence of a significant proportion of zero-lag functional connectivity and indicate that in-antiphase synchrony is preferentially happening along the IH direction, in agreement with previous intracranial observations. However, there are also some discrepancies between these two analyses. For example, the relationship between the type of synchrony (in-phase or in-antiphase) and the orientation of the electrode pairs (IH or AP) was much stronger in the first (average reference) analysis than in the second (SCD). To explain these discrepancies, we hypothesized that they are due to differences in spatial scales: the average reference is known to favor large spatial scales whereas SCD is known to favor smaller spatial scales. In order to disentangle the effect of using potentials measured against a common reference versus current densities on the one hand and the effect of using measures sensitive to different spatial scales, on the other hand, we modified the parameterization of the SCD to make it sensitive to different spatial scales, including a scale similar to the average reference. In Figure 8, we show three types of results: topomaps depicting the phase of coherency using probe electrodes (C6 and Oz), polar plots (as in Figure 3) for in-phase and in-antiphase synchrony, and the orientation index (as in Figure 6^1^). These results are all for 70 Hz, and they are shown for the average reference (as in the first analysis; rightmost columns), a parameterization of SCD that produces signals on a similar spatial scale than the average reference (second column from the right), the SCD with the default parameterization (as in the second analysis; middle column), and two additional SCD parameterization that resolved yet smaller spatial scales. The similarity of the results obtained with the average reference and the SCD parameterized at a similar spatial scale is striking and the contrast between this “large spatial scale SCD” and the “normal SCD” is also very clear. These observations strongly support that the much larger effect sizes found in the first analysis are mostly due to the use of a reference scheme sensitive to differences of potential at a relatively large spatial scale rather than being due to the reference scheme creating “false” zero-lag synchrony.

**Figure 8.**
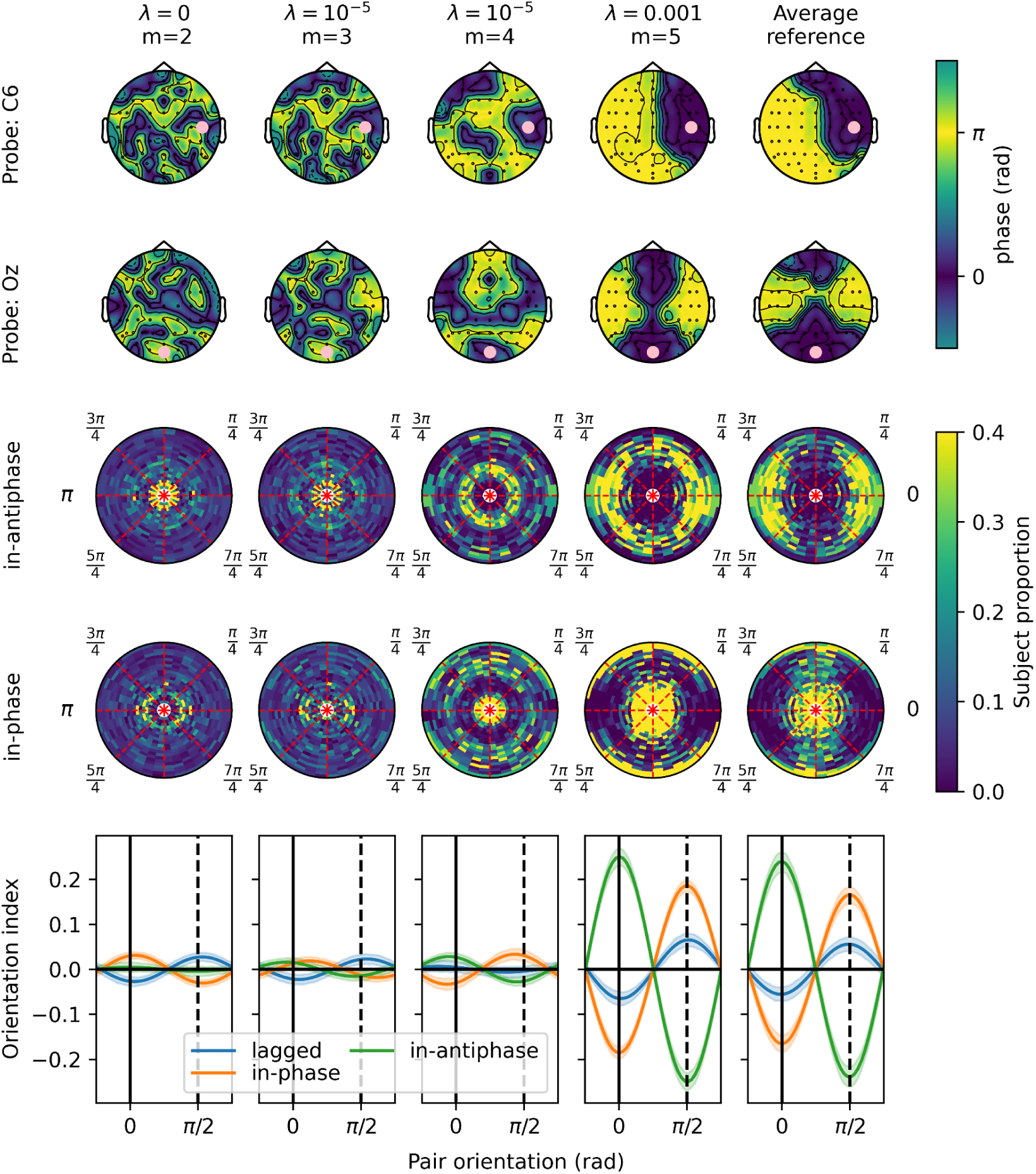
Effect of spatial scales on the property of zero-lag connectivity. The two first rows are showing topomaps of the phase of coherency between a probe electrode (first row: C6; second row: Oz) and the other electrodes. The next two rows show the polar heatmaps of the proportion of subjects showing either in-antiphase (third row) or in-phase (fourth row) synchrony, the r coordinate corresponding to the inter-electrode distance and the angular coordinate corresponding to the orientation of the electrode pairs (0 rad: IH; π/2 rad: AP). The fifth row shows the mean value of the orientation index with the shaded regions indicating the 95% IC across subjects. All computed for the coherency at 70 Hz.

## Discussion

Current approaches for estimating functional connectivity generally reject all zero-lag synchronization as an artifact of volume conduction. Demonstrating that zero-lag connectivity not only exists but is present in significant proportion in EEG recordings would lead to the development of new connectivity measures capable of disentangling true zero-lag connectivity and volume conduction. Such methodological developments are essential to unveil the properties of this type of functional connectivity and may prove to be particularly important in the study of oscillatory circuits and cell assemblies involved in neural binding and in interhemispheric communication. Further, properly disentangling the presence of volume conduction from functional zero-lag connectivity is fundamental in properly estimating the cortical sources responsible for the EEG.

Building on our previous results with intracranial recordings showing a clear presence of zero-lag connectivity predominantly synchronized in antiphase between homotopic regions, here we evaluated if such zero-lag connectivity is present in scalp EEGUnder the *a priori* hypothesis, stemming from our previous results^12^, that we would observe higher in-antiphase synchronization between IH than AP channel pairs, we investigated EEG recordings in 63 subjects re-referenced to a robust average reference. We observed a striking difference in the proportion of in-phase versus in-antiphase synchronization in these two directions, with a clear dominance of in-antiphase synchronization in the IH pairs but not in the AP pairs. These results correspond to our expectations given the zero-lag connectivity observed between homotopic brain regions in intracranial recordings.

Further, in order to demonstrate that these observations cannot be attributed to differences in how volume conduction propagates in the IH and AP directions, we estimated the EEG cortical sources, spatially randomized them and simulated how these sources would propagate through volume conduction to the scalp. All traces of phase preference in the IH versus AP directions disappeared after such randomization, supporting that if such a directionality bias in volume conduction exists, it is small enough not to be statistically significant in our analyses and cannot be the cause of the huge effect size observed in the EEG.

We also addressed the possibility that such synchronization could be due to the use of a common reference. We performed a second analysis using SCD (a reference-free measure of current sources generated by neuronal activity) and further described how patterns of zero-lag synchrony varied as a function of the topology and the frequencies. Although this second analysis confirmed the presence of significant zero-lag connectivity, it also revealed smaller effect sizes and less clear topological patterns. Under the hypothesis that such a discrepancy could be due to these two measures (average reference and SCD) resolving the EEG activity at different spatial scales rather than be caused by the reference injecting false zero-lag synchrony, we performed in third analysis where we modified the spatial filtering properties of the SCD to make it sensitive to spatial scales similar to the average reference. In doing so, we were able to show that SCD thus parameterized produces very similar results to those obtained in the first analysis using an average reference.

Therefore, given 1) the previous demonstration of in-antiphase synchronization between cell assemblies of homotopic brain regions^12^, 2) the very clear pattern of in-antiphase synchronization present specifically in a set of homotopic interhemispheric channel pairs, 3) the confirmation through simulation that such a pattern cannot be ascribed to a directional preference of volume propagation due to the anisotropy of head tissues, and 4) the demonstration that these results cannot be attributed to apparent synchrony generated by the reference scheme, our findings strongly support that zero-lag synchrony has a significant contribution to genuine EEG functional connectivity. Although these zero-lag synchronization patterns are present in EEG at a smaller spatial scale, they are strongest at a larger scale where they show a very clear pattern of interhemispheric in-antiphase connectivity and anteroposterior in-phase connectivity.

Zero-lag connectivity is likely to represent a basic mechanism of communication and synchronization since it has been observed here using baseline activity (not during stimulation and after at least 1 s of non-stimulation) and it has been observed in intracranial recordings for the waking state as well as during rapid-eye movement (REM) and non-REM sleep. As such, its properties are similar to the default-mode network, resting alpha, and sleep slow waves, i.e., basic phenomena that spontaneously emerge when the brain is not otherwise engaged in more demanding tasks and that are potentially involved in neuronal homeostatic processes and general background synchronization of distributed networks. This assumption would be compatible in the case of IH in-antiphase synchrony with IH inhibition being reduced from baseline level during unimanual contraction^24^ and IH inhibition being reduced in musician^31^ (i.e., musician have to learn to “uncouple” both hands for example to allow them to play one melody with the right hand while simultaneously playing a different rhythm with the left hand). Such a hypothesis, however, requires further study to assess how zero-lag connectivity varies in different contexts.

Although we used only the baseline EEG activity for our analyses, it is possible that our results are impacted by long-lasting (i.e., remaining significant even 1s after the end of the stimulus window) entrainments of spatially extended neural networks. This possibility does not impact our main conclusions regarding the presence of zero-lag synchrony in EEG but may impact the reproducibility of the specific patterns of zero-lag synchrony that we reported. Also, IH in-antiphase synchrony was the most prominent when analyzing EEG at large spatial scales whereas our previous results with intracranial recordings (hence, on a much smaller spatial scale) were reporting in-antiphase synchrony very specifically between homotopic brain regions but not with other regions close to their homotopic counterpart. Although both studies are consistent (in-antiphase synchrony in the IH direction), we cannot yet exclude the possibility that these two observations are due to different mechanisms at different spatial scales.

To conclude, our results suggest that the properties of in-phase and in-antiphase zero-lag connectivity need to be determined to accurately assess their contribution to functional connectivity. Such methodological advances are not only important to complement analyses of lagged connectivity, but also to better constrain the underdetermined problem of source estimation in M/EEG. Further, from a clinical point of view, better understanding the relationship between IH inhibition and IH in-antiphase synchrony might be important in devising more effective transcranial magnetic stimulation protocols for stroke recovery^32^ and for conditions showing abnormal information integration and reduced corpus callosum connectivity such as autism spectrum disorder^33, 34^.

## Acknowledgments

This research is supported by the Azrieli Centre for Autism Research (ACAR), the Fonds de recherche du Québec – Santé (FRQS), the UCSD KAVLI Institute Innovative Research Program (grant awarded to JDL), and grant # NIH/NIA RO1 AG18030 awarded to JT. We also want to thank Paul L. Nunez for providing useful comments on this manuscript.

## Online Methods

### Participants

EEG data and T1-weighted scans were collected from 63 healthy adults between 22 and 80 years of age (see Figure 9.a for age and sex distribution). Individuals with psychiatric disorders, a history of neurological illness, any contraindications for magnetic resonance imaging, or uncorrected visual problems were excluded from the study. All participants provided written informed consent.

**Figure 9.**
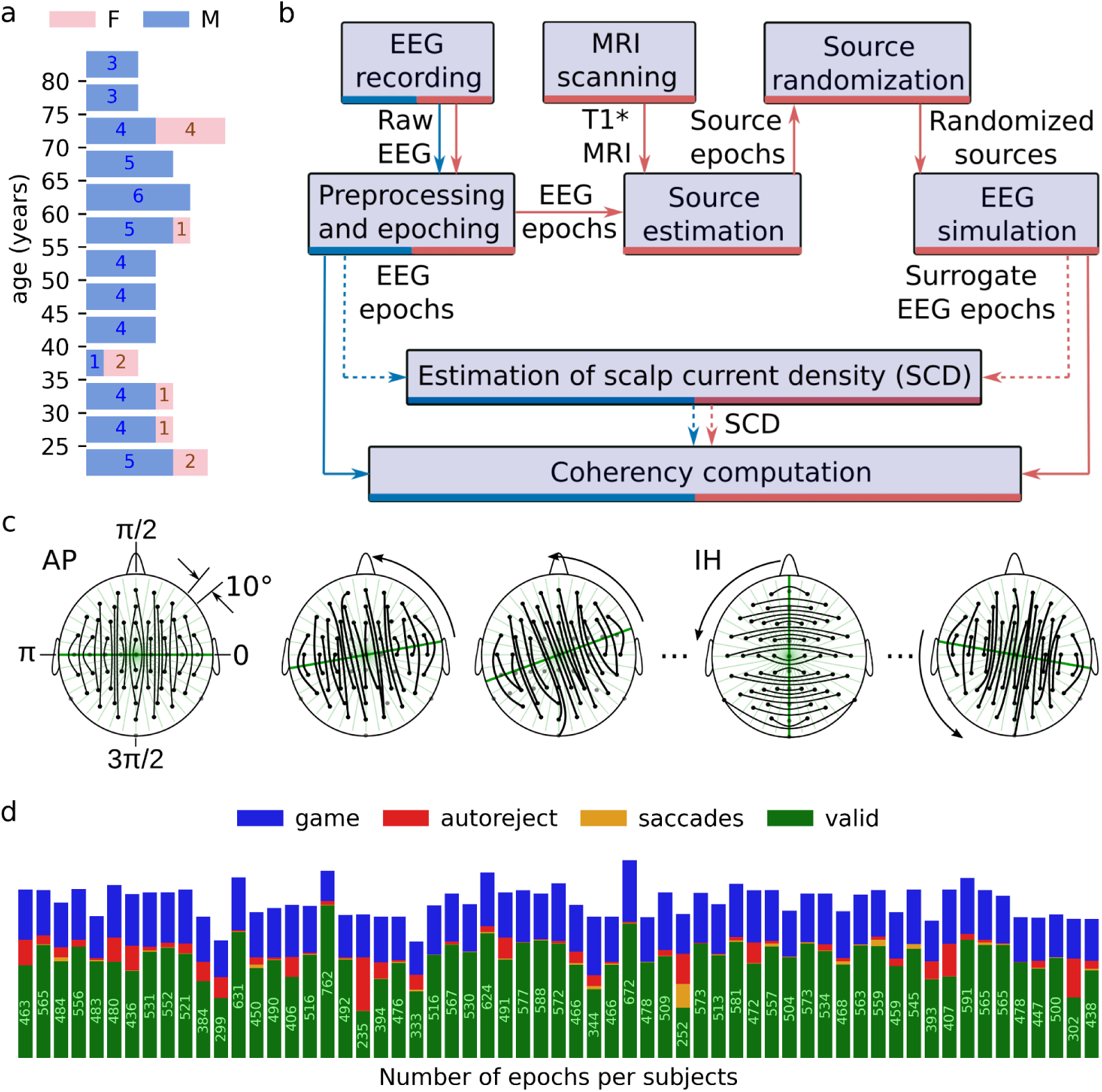
Data collection and preprocessing. a) Age and sex distribution of the sample per 5-year bins (blue: males; pink: females). The numbers in the bars indicate the number of individuals of the corresponding age and sex. b) Diagram showing the processing pipeline for computing coherency from the recorded (blue) and surrogate (red) EEG. Dashed arrows indicate the additional computation of scalp current densities for the second analysis. c) Interhemispheric (IH) and anteroposterior (AP) channel pairs were used in the first analysis. For the following analysis, this concept was generalized as shown in this panel. Channel pairs were selected if they were more or less symmetric to a line passing through Cz and rotated by 10° increment between [0, π[(i.e., IH and AP pairs are obtained when the symmetry axis is at π/2 and 0 rad, respectively). Channels are shown as dots; black curved lines indicate paired channels. The method used to establish this pairing is further described below. d) Number of epochs that are either valid (exact numbers are written in the green bars) or rejected due to game events (described below), statistical outliers (autoreject), and saccades. Each bar represents an individual subject.

### Overview of the analytic pipeline

We implemented an analytic pipeline specifically designed to confirm that genuine zero-lag connectivity (i.e., not due to volume conduction or reference scheme) is entangled amidst volume conduction artifacts. The analytic steps required for this direct demonstration are illustrated by the blue pathway in the schema in Figure 9.b and it comprises EEG acquisition, preprocessing, epoching, and coherency computation.

To investigate if the association between EEG in-phase/in-antiphase synchrony and the orientation of the connections in the transverse plane follows what has been reported in intracranial recordings, we selected IH and AP channel pairs so that the electrodes are as symmetric as possible to a demarcation line taken as the midline channel column (i.e., Cz, Fz, Pz, etc.) for IH and the central channel row (i.e., C3, Cz, C4, etc,) for AP pairs. We generalized this approach for all orientations, by rotating this symmetry line by 10° increments using Cz as the pivot point (Figure 9.b). For each of these increments, we determined for each electrode on one side of the symmetry line which electrode on the other side was positioned the closest to its reflection. To avoid including channel pairs more than once, we considered a given pair only if it was oriented with an angle within 5° of the direction perpendicular to the symmetry plane.

We performed a first analysis using an average mastoid reference re-referenced to a robust average reference and it showed a large proportion of in-antiphase synchronization (see results), specifically in the IH direction. Since the analysis of zero-lag connectivity is suspected to be particularly vulnerable to apparent zero-lag synchrony induced by a common reference, in a second analysis we computed the SCD to eliminate the effect of the reference scheme^35^ and confirm that the results of our first analysis were not due to reference artifacts.

We established a statistical baseline for the effect of volume conduction by using a surrogate dataset. To compute this surrogate dataset, we estimated the cortical sources from the recorded EEG and used these sources to simulate the EEG activity that they would generate, with (surrogate) and without (simulated) randomizing the position of these sources across the cortex (see below for the complete details on the computation of simulated and surrogate signals). As for the recorded EEG, we computed and compared the phase of coherency for the surrogate EEG (red pathway on Figure 9.b). We chose this approach to maximize the “ecological validity” of our surrogate EEG dataset, i.e., all the temporal and spectral features within and between sources used to generate EEG are preserved, only their spatial relationship is randomized.

These different comparisons allow untangling from our estimates of functional connectivity the contribution of 1) volume conduction and 2) zero-lag connectivity. Indeed, in the simulated EEG dataset, the spatial pattern of coherency phases may still be attributed to both directional differences in volume conduction (e.g., due to the shape of the head, to anisotropic conductivity of head tissues, or to the orientation of the dipolar sources) and zero-lag connectivity between cortical EEG sources. However, in the surrogate dataset, only volume conduction contributes to systematic spatial patterns of coherency because the position of the cortical sources has been randomized before simulating the EEG. Therefore, spatial patterns of coherency in the surrogate dataset cannot be ascribed to spatial patterns of functional connectivity between the underlying cortical sources. Consequently, if in our first analysis the in-antiphase synchrony dominates significantly over in-phase synchrony in IH pairs compared to AP pairs in recorded EEG but not in surrogate EEG, we can safely attribute this effect to genuine zero-lag connectivity between homotopic brain regions and conclude that this EEG analysis corroborates previous LFP results.

### EEG acquisition

EEG was recorded using an ActiveTwo system (Biosemi, Inc., Amsterdam, Netherlands) with the Biosemi 10/20 64-electrode cap. The BioSemi ActiveTwo data acquisition system provides amplification at the scalp to minimize movement artifacts. The recording used a Biosemi common-mode sense-driven right leg reference, re-referenced offline to the mastoids and further re-referenced to a robust average reference further down the preprocessing pipeline (see below). Since the experimental protocol used to acquire this dataset was designed with future clinical applications in mind, we took into consideration the fact that the typical placement for EOG beside or directly beneath the eyes was not well tolerated by more sensitive subjects (e.g., autistic children) and placed EOG electrodes on the cheekbones, beneath and lateral to the eyes. Therefore, these two EOG derivations captured simultaneously vertical and horizontal eye movements. For EOG artifact detection, Fp1 and Fp2 channels were also used to supplement the EOG channels.

EEG was recorded during an event-related paradigm in which different combinations of horizontal and vertical drifting sinusoidal gratings were presented in each hemifield. Recordings were divided into 4 s epochs, centered on the stimuli, i.e. 1 s pre-stimulus baseline, 2 s of drifting grating(s) presentation, and 1 s post-stimulus. A game component was also added to this protocol to keep the participant focused on a central fixation point. It required the participants to hit a button as fast as possible upon a subtle change of color of a spaceship centered on the screen to shield themselves from a missile attack. The game was designed to force participants to keep the central fixation point on their central fovea where the color-sensitive cones are concentrated. Game events were recorded and used to reject corresponding epochs. A more thorough description of the game and the experimental paradigm is provided in Lewis et al. (in review).

Since the presence of zero-lag connectivity between homotopic brain regions has been demonstrated irrespective of consciousness states (across sleep stages and in wakefulness) in LFP recordings^12^, this type of functional connectivity is expected to be a fundamental mechanism present regardless of the presence or not of a stimulus. Also, our preliminary analyses showed no clear impact of the stimuli on the patterns we report in this paper. Therefore, we used only the 1 s pre-stimulus baseline period from valid epochs (i.e., without artifacts or game events) for our final analyses, i.e., we do not report on stimulus- or task-related properties of the brain activity^2^ in this paper. We also do not report on demographic (i.e., age and sex) effects because this study aims at demonstrating the existence of a basic mechanism that is expected to be present during the normal operation of a typical brain. The potential effect of tasks, demographic properties, or health-related issues on such zero-lag connectivity can be assessed in follow-up studies, once the existence and significance of EEG zero-lag connectivity are demonstrated by the current study.

### EEG preprocessing and epoching

PyPREP, the Python wrapper for the PREP pipeline^36^, was used to compute a robust average reference, remove line noise, and identify bad channels for later interpolation using MNE-Python^37^. Blinks were automatically detected using a robust peak-to-peak thresholding approach and EEG signals were corrected using a signal-space projection operator relying on manually selected artifact components. A semi-automated approach was used to detect and remove epochs containing lateral saccades. The detection of blinks and saccades was reviewed manually to ensure that it was appropriate for all subjects.

All epochs overlapping a game event were rejected. Autoreject^38^ was further used to clean the final epoched dataset, interpolating or rejecting channels and epochs depending on their level of noise. Figure 9.d shows the number of valid epochs per subject, as well as the number of epochs rejected at these different preprocessing stages.

### MRI acquisition and processing

3D T1-weighted images (Fast Gradient Echo, SPGR; TE = 3.1ms; flip angle = 12°; NEX = 1; FOV = 25cm; matrix=256×256) were recorded in all subjects on a GE Signa EXCITE 3.0T short bore scanner with an eight-channel array head coil. These T1 volumes were processed with CIVET (version 2.1.0; 2016), a fully automated structural image analysis pipeline. CIVET corrects intensity non-uniformities by non-parametric non-uniform intensity normalization (N3)^39^; aligns the input volumes to the Talairach-like ICBM-152-nl template with an affine transformation followed by a nonlinear transformation^40^; classifies the input volumes into white matter, gray matter, cerebrospinal fluid, and background^41^; and extracts the white matter and pial surfaces^42^.

### Source estimation

Cortical sources have been estimated with MNE-Python^37, 43^. To compute a boundary element method (BEM) conductor model of the subjects’ head, CIVET cortical white matter and pial surfaces were imported and converted to FreeSurfer^44^ surfaces. CIVET preprocessed T1w volumes were converted to .mgz using NiBabel^45^, their intensity was normalized and the FreeSurfer watershed algorithm was used to extract the surface of the scalp, and both the inside and the outside of the skull. To further constrain the watershed algorithm and ensure compatibility between the inner skull and cortical surfaces, Trimesh^46^ and NiBabel were used to fill and voxelize the volume enclosed by the pial surface and save it as a brain mask. Further custom scripts were used to fix any topological defects in the meshes and ensure watertightness and non-intersecting surfaces. BEM surfaces were further decimated to 5120 triangles based on the FreeSurfer spherical co-registration algorithm.

Our source models included about 8200 sources (varies slightly from subject to subject) placed at the position of randomly selected cortical mesh vertices and oriented normal to this surface (i.e., perpendicular to the cortex). For source estimation, we used the eLORETA^47^ inverse operator and a diagonal noise covariance matrix to avoid cancelling-out possible contributions of zero-lag connectivity to non-diagonal covariance components.

The quality of the co-registration of the head shape, the electrodes, and the dipolar sources is illustrated for an example subject^3^ in Figure 10.a-c. We also validated the properties of the source reconstruction leadfield, which is the matrix that encodes how source activity propagates to scalp sensors through volume conduction. Since the orientation of these dipoles is fixed perpendicular to the cortex, each dipolar source can be represented by its scalar amplitude. The leadfield is therefore a matrix of size N X M, with N being the number of channels and M being the number of dipoles. By computing the N X N correlation matrix associated with this leadfield, we obtain for every pair of channels a Pearson’s coefficient of correlation that indicates how much these two channels depend on the same sources and, therefore, how much they are likely to show in-phase (positive correlation) or in-antiphase (negative correlation) synchrony due to the activity of these sources. Although the full correlation matrix is difficult to interpret, the leadfield can be validated by visualizing individual rows of this matrix as topographical maps (topomaps) describing the correlation between a specific channel (i.e., a specific row of the correlation matrix) used as a probe and the other channels. Using this approach, we can see that the leadfield creates a high correlation between nearby electrodes and anti-correlation at longer distances (Figure 10.d), as expected from electrical fields generated by dipolar sources. Alternatively, we can also plot these correlations as topomaps for IH and AP pairs of channels (Figure 10.e). From these plots, we can see that the properties of the leadfield are not significantly different in the IH versus the AP directions, which already suggests that the volume conduction is unlikely to be responsible for systematic differences in phase synchrony depending on these directions.

**Figure 10.**
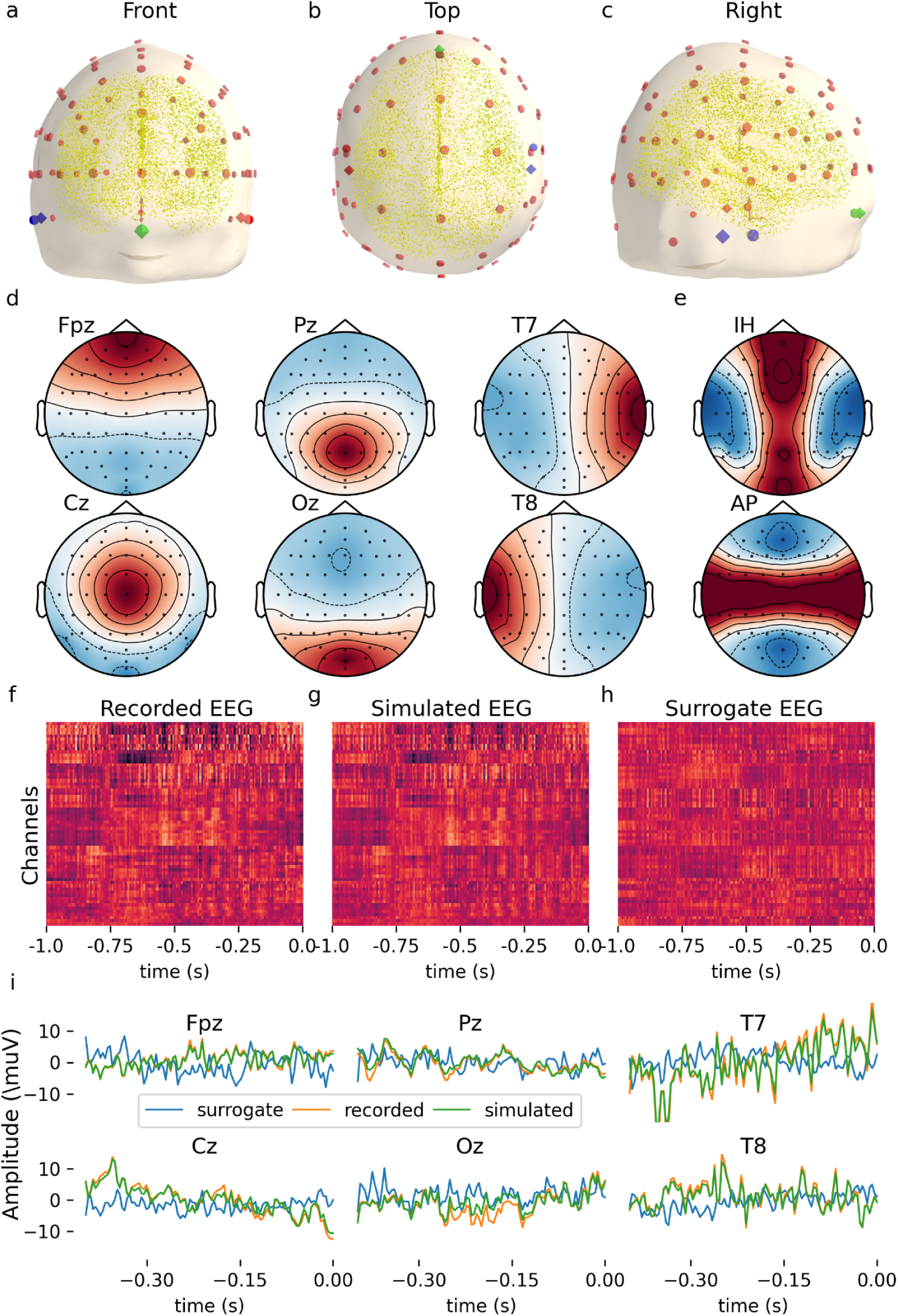
Validation of source estimation and EEG simulation. All results are illustrated for an example subject and panels f-h show results computed for the first valid epoch of this subject. a-c) Rendering of the head model used for source estimation and EEG simulation, including the scalp surface extracted from the MRI, EEG electrodes (red disks), fiducials (red: left preauricular, green: nasion, bleu: right preauricular; spheres: MRI fiducials, octahedron: EEG montage fiducials), and dipolar sources (small yellow spheres). d) Topomaps showing the correlation of leadfield components between probe channels (identified over the plots) and the other channels. e) Similar as in d, but showing the correlations between the IH and AP pairs. f-h) Recorded (f), simulated (g), and surrogate (h) EEG signals are shown as a heatmap for all 64 channels. i) EEG signals are shown in panels f-h overlaid, for six examples of channels. The amplitude of surrogate signals in (h, i) has been multiplied by a factor of 10 for them to be on the same scale as the recorded and simulated EEG. Surrogates EEG signals are generally weaker because the cortical sources are by design less spatially correlated and, therefore, their vectorial summation produces smaller EEG potentials. Such a homogeneous scaling factor has no impact on coherency.

### Functional connectivity

The coherency between two signals *x* and *y* can be defined as a function of frequency *f* as

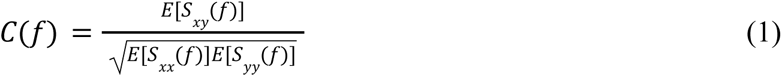

In this equation, E[] indicates the average across epochs and *S_xy_* stands for the cross-spectral density which can be defined with respect to the cross-correlation function between *x* and *y*

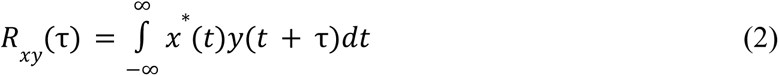

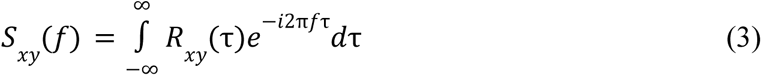

with *x*\* being the complex conjugate of *x* which is equal to *x* in our case since EEG signals are real-valued. These coherency values have been computed between every channel pair using Python-MNE.

Note that (1) is slightly different from the definition of coherence based only on the absolute value of the coherency and therefore taking only real values. In contradistinction, coherency takes complex values of which the modulus indicates the coherence and the phase indicates the phase difference between the spectral content of the two signals at the corresponding frequency (i.e., the phased difference of the two band-passed signals).

The phase of the estimated coherency can be obtained using

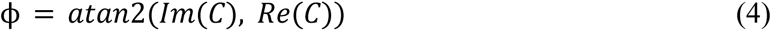

Further, to avoid issues due to the cyclic nature of phases, average phases were computed using

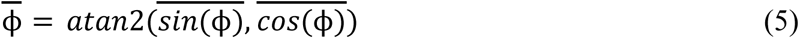

where 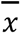 indicates the average of x.

### Determining in-phase and in-antiphase synchrony

In-phase and in-antiphase synchrony is determined using a threshold T. Channel pairs with phases ϕ such that |ϕ| < T or |ϕ-π| < T are considered as in-phase or in-antiphase, respectively. Unless specified otherwise, T=0.05 has been used in our analyses. This corresponds to 0.8% of a cycle. We observed that increasing T was generally increasing the effect sizes, but larger T values are less specific and include connections that might be better characterized as having small but non-zero delays. The value 0.05 was used as a tradeoff between these two objectives (maximizing effect sizes while ensuring that the selected connections are representative of zero-lag synchrony). The effect of this threshold is further explored in Supp. Figure 10.

### Orientation index

To determine if in-phase and in-antiphase synchronization is preferentially oriented in a given direction, we defined an orientation index. For a given set of amplitudes (Y) and angles (Θ), we define this index by first demeaning and normalizing Y to a unitary standard deviation:

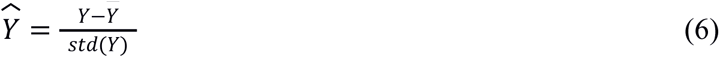

Then, the orientation index for a given orientation θ is defined as follows

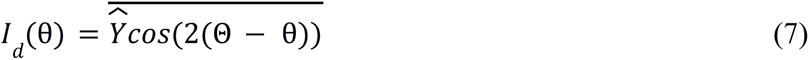

In short, this index is a weighted average of the normalized amplitude, using as weight a sinusoidal function that peaks at any π multiple of the target angle θ. For example, *I_d_* (0) will be close to 1 if all the values in Θ are close to 0 or π (IH pairs) and it will be close to -1 if all the values in Θ are close to π/2 or 3π/2 (AP pairs). The inverse is true for *I_d_*(π/2).

### Zero-lag index

We also defined a zero-lag index to compare the dominance of either in-phase or in-antiphase synchronization. To define this index, we first compute a probability density function *f*(Φ)for a given set of phase estimatesΦusing a kernel density estimation after restraining the values of Φin the [− π/2, 3π/2[interval, using((Φ − π/2) % 2π) + π/2with % being the modulo operator.

We then consider

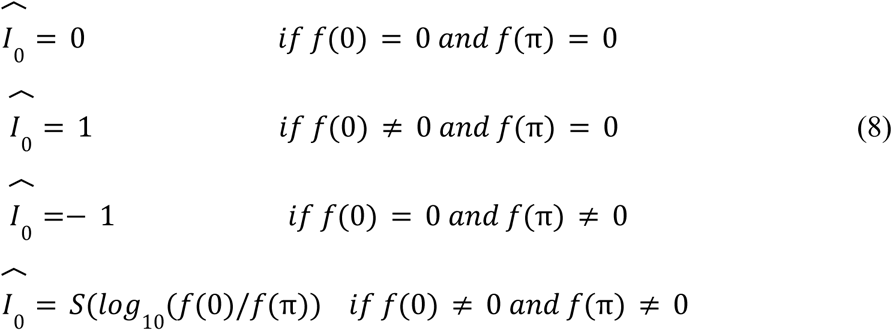

with

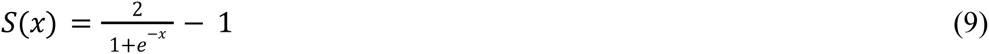

being a sigmoidal function smoothly varying from -1 to 1 as x varies from -∞ to +∞ and crossing the origin. We finally obtain the zero-lag index by normalizing 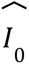 to ensure that its values go toward zero as the distribution of the phases moves away from either at 0 or π radian:

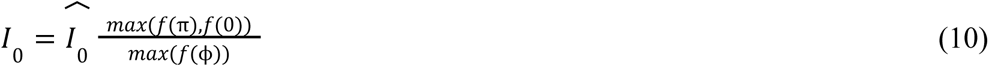

This index is similar to what has been previously proposed^12^ but it has been further modified to take into account some situations where the original index was undetermined, to normalize it to [-1, 1] interval, and to weigh it so that it takes values close to 1 or -1 only when the peak of the distribution is at 0 (in-phase) or π(in-antiphase) radian.

### EEG simulation

EEG simulation was performed with MNE-Python using the estimated source time-series and the same forward models that were used for source estimation. This means that if the processes of source estimation and EEG simulation were invertible, the simulated EEG would be identical to the recorded EEG. Unfortunately, the source estimation process is an under-determined problem and is therefore not invertible. Thus, although the goal of source estimation is to fit the source signals in such a way that once combined with the forward model, the difference between the simulated EEG and the recorded EEG is minimal, this difference is never null in practice. We, therefore, validated the precision of the EEG simulations and confirmed that the EEG recordings are accurately reproduced by simulating the EEG from the estimated sources (Figure 10.f-i). The same simulation process was used to generate the surrogate EEG dataset, except that the order of the sources was randomly swapped in the source matrix (i.e., the association between the sources positions and the time-series describing their activity was randomized) to break the relationship between the sources functional connectivity and their location in the cortex. This means that any systematic spatial pattern of in-phase and in-antiphase synchronization between EEG channels on the surrogate dataset would be due to volume conduction only since the component due to source functional connectivity has been spatially randomized.

### Scalp current density

For our second analysis, scalp current densities (also known as current source density (CSD)^4^ and surface Laplacian) have been computed using MNE-Python, which ported to Python the Matlab CSD Toolbox^48^. This implementation uses spherical splines to compute the curvature of the electrical field and this process depends on two parameters adjusting the spline stiffness (m) and the solution smoothness (λ). For this analysis, we used the default MNE-Python values for these parameters: m=4 (validated in the original publications on this approach to SCD estimation^35, 49^) and λ=10^-5^. However, since these parameters can be used to modify the spatial filtering properties of the SCD, in a third analysis we varied these values to compare the results of our first analysis using a reference average (known to favor low spatial frequency^50^) with those of our second analysis using SCD (which is known to favor high spatial frequency^50^ with the default parameterization described above). By parameterizing the SCD to have a similar spatial sensitivity than the average reference, we show that the differences between our first two analyses are mostly due to this difference in spatial scales and not to zero-lag synchrony between channels being introduced by the use of a common reference.

It is worth noting that aside from being independent from any reference, SCD helps reduce the effect of volume conduction. It is expected to be dominated by contributions from radial dipoles in superficial gyral surfaces of the neocortex so, as opposed to the average reference, it tends to emphasize superficial and localized sources and to deemphasize deep sources and widespread coherent superficial sources^50^.

### Code availability

All signal processing and statistical analyses were performed in Python using open-source software. The scripts written for these analyses are available on the first author GitHub page (https://github.com/christian-oreilly).

## Supporting documents

**Supp. Figure 1.**
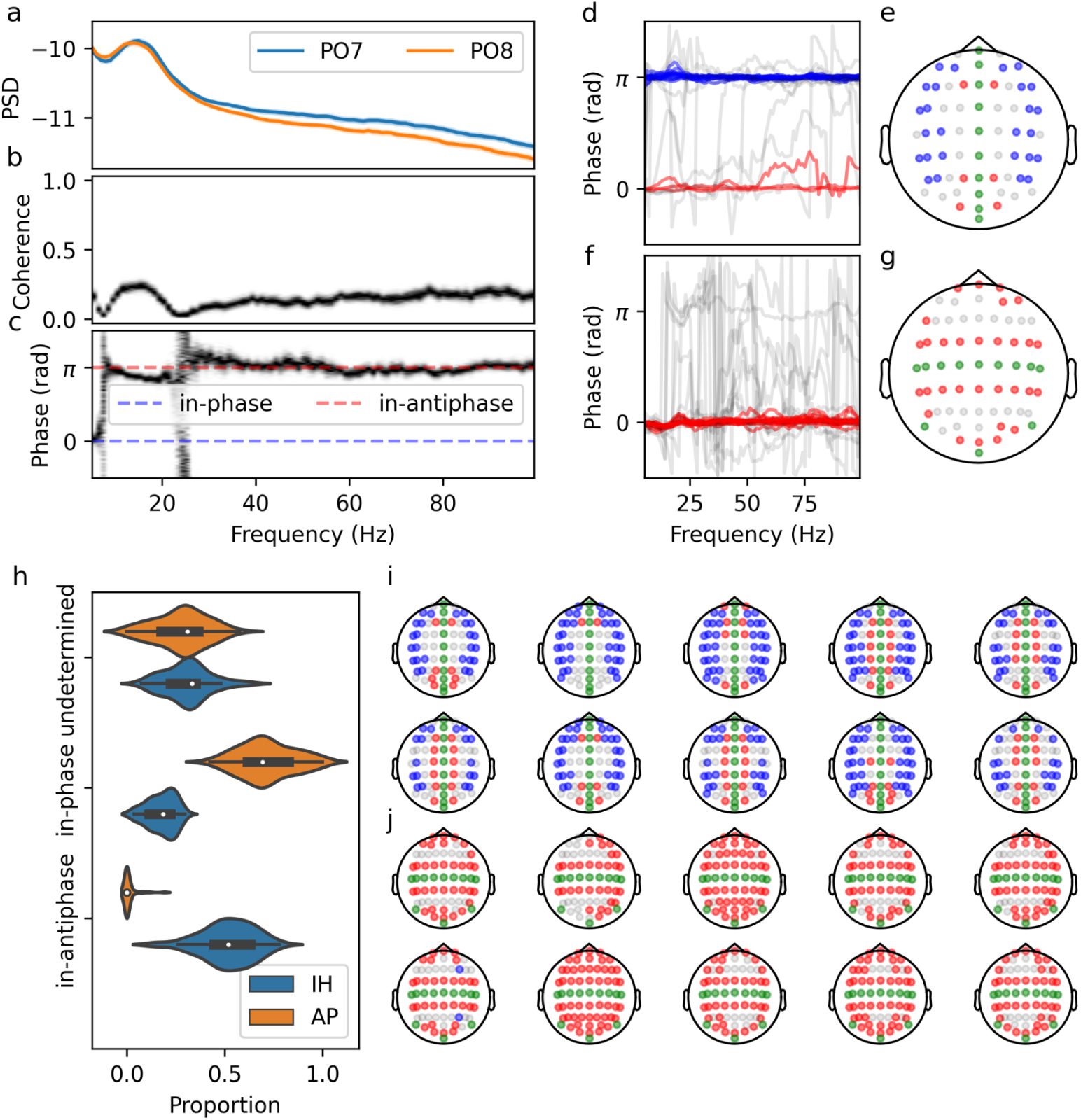
Comparative analysis of IH and AP channel pairs for EEG signals simulated from estimated sources. a-c) Example of PSD (a), coherence (b), and coherency phase (c) for the (PO7, PO8) channel pair on an example subject. d-g) These panels show the same data as for Figure 3.d-g, but for the simulated EEG. h) Distribution of the proportion of pairs per synchronization type, across subjects (same as for Figure 3.j). i,j) Color-coded dominant coherency phase (blue: in-antiphase; red: in-phase; grey: undetermined; green: unpaired) for 10 randomly selected subjects, for the IH (i) and the AP (j) directions.

**Supp. Figure 2.**
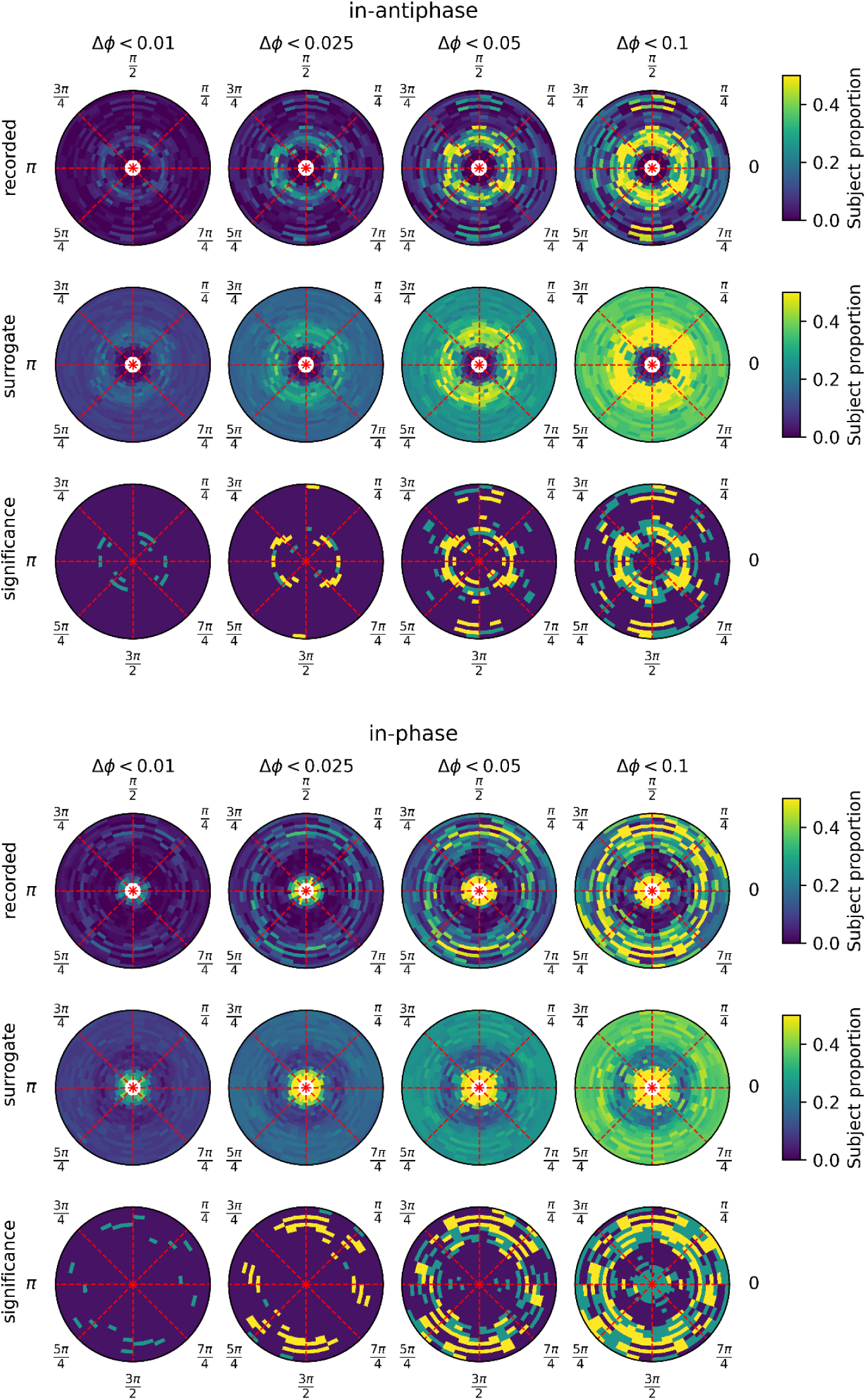
Polar plots showing the proportion of subjects with in-antiphase (upper panel) or in-phase (bottom panel) synchrony as a function of the channel pair orientation (angular coordinate) and the inter-electrode distance (r coordinate). Each column shows the results for a different threshold T used to classify channel pairs as in-phase or in-antiphase (0.01, 0.025, 0.05, 0.1). On the polar plots, a 0 rad angle corresponds to IH, whereas π/2 rad corresponds to AP. The first row of each panel gives the values from the recording, the second row shows the 99th percentile computed on the surrogate distribution, and the third row shows the statistical significance (dark blue: not statistically significant; light blue: significant at p<0.01, not corrected for multiple tests; yellow: significant at p<0.01, Bonferonni corrected for testing independently for 834 channel pairs). For this analysis, the estimated phases of the coherency have been computed for 70 Hz.

**Supp. Figure 3.**
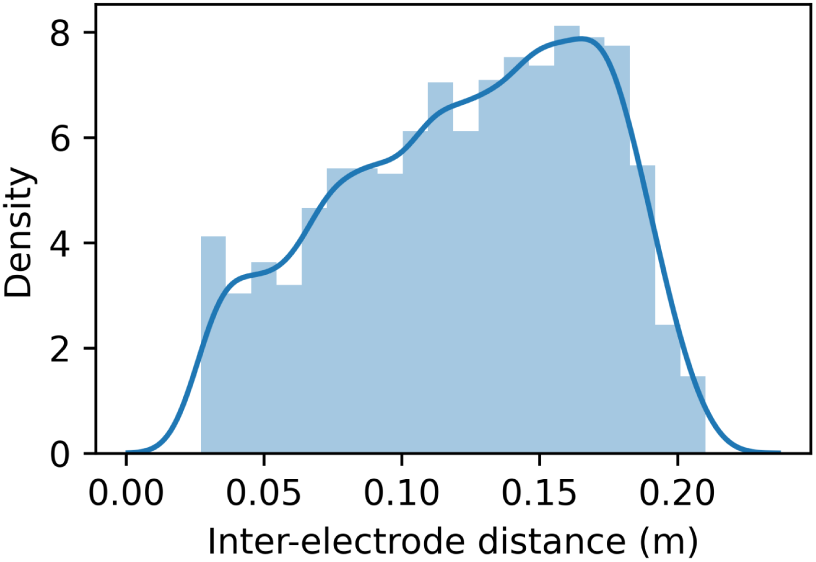
Distribution of the inter-electrode distances.

## Footnotes

1 Except that the orientation index is computed using boolean values equal to one if the pairs are classified as the corresponding type of synchrony instead of according to thresholds computed from a surrogate dataset. This is done for computational reasons (i.e., to avoid the expensive computation of surrogate datasets for all these conditions) and because this less demanding approach is sufficient for comparing between these different conditions. This, however, explains why the orientation index in the panel in the fifth row, third column of Figure 8 has smaller amplitude than the corresponding plot in the first row, fourth column of Figure 6.

2 Strictly speaking, these recordings are still done during a game protocol and some background stimulation that are not time-locked — like a moving starfield — is still present. Also, although the epochs considered cover a period starting 1 s after the end of the last stimulus period, some residual activation may still be present. However, we do not see this as a limitation since the goal of this study is not to associate EEG zero-lag connectivity with a specific paradigm (e.g., event-related or resting-state) but to demonstrate that such instantaneous activity exists and that it is a significant source of functional connectivity in EEG.

3 To allow comparison, we use the same participant for examples across the paper.

4 The name current source density is arguably more often used than scalp current density, but we prefer the later name in this context since “current source” can easily be confused with the cortical sources estimated by inverse modeling.

## References

1. Nunez, P. L. et al. EEG coherency. I: Statistics, reference electrode, volume conduction, Laplacians, cortical imaging, and interpretation at multiple scales. Electroencephalogr. Clin. Neurophysiol. 103, 499–515 (1997).

2. Plonsey, R. & Heppner, D. B. Considerations of quasi-stationarity in electrophysiological systems. Bull. Math. Biophys. 29, 657–664 (1967).

3. Brookes, M. J., Woolrich, M. W. & Barnes, G. R. Measuring functional connectivity in MEG: A multivariate approach insensitive to linear source leakage. NeuroImage 63, 910–920 (2012).

4. Colclough, G. L., Brookes, M. J., Smith, S. M. & Woolrich, M. W. A symmetric multivariate leakage correction for MEG connectomes. Neuroimage 117, 439–448 (2015).

5. Nolte, G. et al. Identifying true brain interaction from EEG data using the imaginary part of coherency. Clin. Neurophysiol. Off. J. Int. Fed. Clin. Neurophysiol. 115, 2292–2307 (2004).

6. Stam, C. J., Nolte, G. & Daffertshofer, A. Phase lag index: assessment of functional connectivity from multi channel EEG and MEG with diminished bias from common sources. Hum. Brain Mapp. 28, 1178–1193 (2007).

7. Varela, F., Lachaux, J. P., Rodriguez, E. & Martinerie, J. The brainweb: phase synchronization and large-scale integration. Nat. Rev. Neurosci. 2, 229–239 (2001).

8. Campo, A. T. et al. Feed-forward information and zero-lag synchronization in the sensory thalamocortical circuit are modulated during stimulus perception. Proc. Natl. Acad. Sci. 116, 7513–7522 (2019).

9. Engel, A. K., König, P., Kreiter, A. K. & Singer, W. Interhemispheric Synchronization of Oscillatory Neuronal Responses in Cat Visual Cortex. Science 252, 1177–1179 (1991).

10. Gray, C. M., König, P., Engel, A. K. & Singer, W. Oscillatory responses in cat visual cortex exhibit inter-columnar synchronization which reflects global stimulus properties. Nature 338, 334–337 (1989).

11. Nikouline, V. V., Linkenkaer-Hansen, K., Huttunen, J. & Ilmoniemi, R. J. Interhemispheric phase synchrony and amplitude correlation of spontaneous beta oscillations in human subjects: a magnetoencephalographic study. NeuroReport 12, 2487–2491 (2001).

12. O’Reilly, C. & Elsabbagh, M. Intracranial recordings reveal ubiquitous in-phase and in-antiphase functional connectivity between homotopic brain regions in humans. J. Neurosci. Res. (2020) doi:10.1002/jnr.24748.

13. Roelfsema, P. R., Engel, A. K., König, P. & Singer, W. Visuomotor integration is associated with zero time-lag synchronization among cortical areas. Nature 385, 157–161 (1997).

14. Dalla Porta, L. et al. Exploring the Phase-Locking Mechanisms Yielding Delayed and Anticipated Synchronization in Neuronal Circuits. Front. Syst. Neurosci. 13, (2019).

15. Vicente, R., Gollo, L. L., Mirasso, C. R., Fischer, I. & Pipa, G. Dynamical relaying can yield zero time lag neuronal synchrony despite long conduction delays. Proc. Natl. Acad. Sci. U. S. A. 105, 17157–17162 (2008).

16. Viriyopase, A., Bojak, I., Zeitler, M. & Gielen, S. When Long-Range Zero-Lag Synchronization is Feasible in Cortical Networks. Front. Comput. Neurosci. 6, (2012).

17. Bullock, T. H. et al. EEG coherence has structure in the millimeter domain: subdural and hippocampal recordings from epileptic patients. Electroencephalogr. Clin. Neurophysiol. 95, 161–177 (1995).

18. Bullock, T. H. et al. Temporal fluctuations in coherence of brain waves. Proc. Natl. Acad. Sci. U. S. A. 92, 11568–11572 (1995).

19. Palmer, L. M. et al. The Cellular Basis of GABAB-Mediated Interhemispheric Inhibition. Science 335, 989–993 (2012).

20. Carson, A. M. Neural correlates of the extreme male brain theory in adolescents with and without autism spectrum disorders. (2014).

21. Ni, Z. et al. Two Phases of Interhemispheric Inhibition between Motor Related Cortical Areas and the Primary Motor Cortex in Human. Cereb. Cortex 19, 1654–1665 (2009).

22. Gerloff, C. & Andres, F. G. Bimanual coordination and interhemispheric interaction. Acta Psychol. (Amst.) 110, 161–186 (2002).

23. Geffen, G. M., Jones, D. L. & Geffen, L. B. Interhemispheric control of manual motor activity. Behav. Brain Res. 64, 131–140 (1994).

24. Nelson, A. J., Hoque, T., Gunraj, C., Ni, Z. & Chen, R. Bi-directional interhemispheric inhibition during unimanual sustained contractions. BMC Neurosci. 10, 31 (2009).

25. Cazzoli, D., Wurtz, P., Müri, R. M., Hess, C. W. & Nyffeler, T. Interhemispheric balance of overt attention: a theta burst stimulation study. Eur. J. Neurosci. 29, 1271–1276 (2009).

26. Hilgetag, C. C., Théoret, H. & Pascual-Leone, A. Enhanced visual spatial attention ipsilateral to rTMS-induced ‘virtual lesions’ of human parietal cortex. Nat. Neurosci. 4, 953–957 (2001).

27. Forss, N., Hietanen, M., Salonen, O. & Hari, R. Modified activation of somatosensory cortical network in patients with right-hemisphere stroke. Brain 122, 1889–1899 (1999).

28. Seyal, M., Ro, T. & Rafal, R. Increased sensitivity to ipsilateral cutaneous stimuli following transcranial magnetic stimulation of the parietal lobe. Ann. Neurol. 38, 264–267 (1995).

29. Cohen, J. Statistical power analysis for the behavioral sciences. (L. Erlbaum Associates, 1988).

30. Sawilowsky, S. New Effect Size Rules of Thumb. J. Mod. Appl. Stat. Methods 8, (2009).

31. Ridding, M. C., Brouwer, B. & Nordstrom, M. A. Reduced interhemispheric inhibition in musicians. Exp. Brain Res. 133, 249–253 (2000).

32. Boddington, L. J. & Reynolds, J. N. J. Targeting interhemispheric inhibition with neuromodulation to enhance stroke rehabilitation. Brain Stimulat. 10, 214–222 (2017).

33. Travers, B. G. et al. Atypical development of white matter microstructure of the corpus callosum in males with autism: a longitudinal investigation. Mol. Autism 6, 15 (2015).

34. Oberman, L. M. et al. Transcranial magnetic stimulation in autism spectrum disorder: Challenges, promise, and roadmap for future research. Autism Res. Off. J. Int. Soc. Autism Res. 9, 184–203 (2016).

## References

35. Perrin, F., Bertrand, O. & Pernier, J. Scalp Current Density Mapping: Value and Estimation from Potential Data. IEEE Trans. Biomed. Eng. **BME****-**34, 283–288 (1987).

36. Bigdely-Shamlo, N., Mullen, T., Kothe, C., Su, K.-M. & Robbins, K. A. The PREP pipeline: standardized preprocessing for large-scale EEG analysis. Front. Neuroinformatics 9, (2015).

37. Gramfort, A. et al. MNE software for processing MEG and EEG data. NeuroImage 86, 446–460 (2014).

38. Jas, M., Engemann, D. A., Bekhti, Y., Raimondo, F. & Gramfort, A. Autoreject: Automated artifact rejection for MEG and EEG data. NeuroImage 159, 417–429 (2017).

39. Sled, J. G., Zijdenbos, A. P. & Evans, A. C. A nonparametric method for automatic correction of intensity nonuniformity in MRI data. IEEE Trans. Med. Imaging 17, 87–97 (1998).

40. Collins, D. L., Neelin, P., Peters, T. M. & Evans, A. C. Automatic 3D intersubject registration of MR volumetric data in standardized Talairach space. J. Comput. Assist. Tomogr. 18, 192–205 (1994).

41. Zijdenbos, A. P., Forghani, R. & Evans, A. C. Automatic ‘pipeline’ analysis of 3-D MRI data for clinical trials: application to multiple sclerosis. IEEE Trans. Med. Imaging 21, 1280–1291 (2002).

42. Kim, J. S. et al. Automated 3-D extraction and evaluation of the inner and outer cortical surfaces using a Laplacian map and partial volume effect classification. NeuroImage 27, 210–221 (2005).

43. Gramfort, A. et al. MEG and EEG data analysis with MNE-Python. Front. Neurosci. 7, (2013).

44. Fischl, B. FreeSurfer. NeuroImage 62, 774–781 (2012).

45. Brett, M. et al. nipy/nibabel: 3.0.0. (Zenodo, 2019). doi:10.5281/zenodo.3583002.

46. Dawson-Haggerty et al. trimesh. (2020).

47. Pascual-Marqui, R. D., Michel, C. M. & Lehmann, D. Low resolution electromagnetic tomography: a new method for localizing electrical activity in the brain. Int. J. Psychophysiol. Off. J. Int. Organ. Psychophysiol. 18, 49–65 (1994).

48. Kayser, J. & Tenke, C. E. Principal components analysis of Laplacian waveforms as a generic method for identifying ERP generator patterns: I. Evaluation with auditory oddball tasks. Clin. Neurophysiol. Off. J. Int. Fed. Clin. Neurophysiol. 117, 348–368 (2006).

49. Spherical splines for scalp potential and current density mapping. Electroencephalogr. Clin. Neurophysiol. 72, 184–187 (1989).

50. Nunez, P. L. & Srinivasan, R. Electric Fields of the Brain: The neurophysics of EEG. Electric Fields of the Brain (Oxford University Press, 2006).

